# Inferring a novel insecticide resistance metric and exposure variability in mosquito bioassays across Africa

**DOI:** 10.1101/2024.03.09.584248

**Authors:** Adrian Denz, Mara D. Kont, Antoine Sanou, Thomas S. Churcher, Ben Lambert

## Abstract

Malaria claims approximately 500,000 lives each year, and insecticide-treated nets (ITNs), which kill mosquitoes that transmit the disease, remain the most effective intervention. However, resistance to pyrethroids, the primary insecticide class used in ITNs, has risen dramatically in Africa, making it difficult to assess the current public health impact of pyrethroid-ITNs. Past work has modelled the relation between pyrethroid susceptibility measured in discriminating-dose susceptibility bioassays and ITN effectiveness in experimental hut trials. Here, we introduce a new predictive approach that accounts for heterogeneity in insecticide resistance within wild mosquito populations, for example, due to genetic variability, by incorporating data from newly recommended intensity-dose susceptibility bioassays. We fit our mathematical model to a comprehensive data set that combines discriminating dose bioassays from all over Africa, intensity dose bioassays from Burkina Faso, and concurrent experimental hut trials. Our analysis estimates location- and insecticide-specific variation in resistance heterogeneity in Burkina Faso and quantifies differences in insecticide exposure in bioassays and experimental huts. By providing a mechanistic understanding of these experimental data, our approach could be integrated into malaria transmission models to account for the public health impact of insecticide resistance detected by surveillance programmes.

**Author summary:** Bednets treated with insecticides that kill mosquitoes have been responsible for major reductions in malaria burden over recent decades. However, the spread of insecticide resistance in mosquito populations threatens to slow or reverse these gains. It is therefore important to be able to gauge insecticide resistance in local mosquito populations and the remaining effectiveness of bednets. Important tools for quantifying insecticide resistance include susceptibility bioassays, which expose mosquitoes to specific insecticide doses and measure mortality, and experimental hut trials, which aim to mimic how mosquitoes interact with insecticides on bednets in the field. Here, we develop a mathematical model that incorporates mechanistic features of how mosquitoes interact with insecticides in both types of experiments. We show that this model can accurately predict mosquito mortality in experimental hut trials using data from intensity-dose susceptibility bioassays. These bioassays are more sensitive than traditional discriminating-dose assays while being less costly and logistically demanding than experimental hut trials. Our model provides a more granular understanding of insecticide resistance measurements and could be embedded into malaria transmission models to predict the public health impact of insecticide resistance measured by surveillance programmes.

## Introduction

Malaria kills more than half a million people worldwide each year [1] and is transmitted by mosquitoes of the Genus *Anopheles*. Insecticide treated bednets (ITNs) protect individuals sleeping under them from blood-feeding mosquitoes but more importantly kill mosquitoes, which provides a community-level protection against disease [2–7]. From 2000 until the mid-2010s, the number of deaths from malaria fell dramatically, primarily due to the widespread use of ITNs [8]. However, the emerging resistance in mosquitoes to the pyrethroid insecticides used in ITN formulations has led to a debate about the continued effectiveness of ITNs [9–16]. Large trials were conducted to assess the impact of resistance to pyrethroids on the effectiveness of ITN-based malaria vector control [17, 18], but quantifying the impact of resistance by comparison across epidemiological settings is difficult due to many interfering factors [19] and so the overall impact of mosquito resistance threat remains unclear [20].

In addition to quantifying the malaria burden due to loss of ITN effectiveness through the spread of pyrethroid resistance, it is important to factor in resistance in localised malaria prevention planning. Simulation models of malaria epidemiology [21–23] take into account many epidemiological factors and are increasingly used for the fine-scale planning of intervention strategies. These models require parameterising the differential effects on malaria transmission for each ITN product taking into account insecticide resistance in mosquitoes. In particular, decisions about what ITN product to use in a specific setting depend on its ability to kill mosquitoes with different resistance profiles [24].

Experimental hut trials (EHT) are a standard assay to evaluate ITN products under controlled field conditions [25] and are the primary data source for parameterising their entomological impact in models of malaria epidemiology [13, 26–28]. An experimental hut mimics typical housing in the study area but is specially constructed to retain all entering mosquitoes, with three main hut designs used in Sub-Saharan Africa: East African [29], West African [30] and Ifakara design [31]; which differ in catching efficiency [32]. During an EHT, a human volunteer sleeps inside the hut under either an untreated control net or an ITN, and after each study night, all dead and alive mosquitoes are collected from the hut and their blood-feeding status recorded. From these detailed data, ITN effects on mosquito survival and blood-feeding can be estimated [13, 26–28]. EHTs are relatively laborious and expensive, and so are conducted in a limited number of locations only.

Programmes to monitor pyrethroid resistance occurrence across Sub-Saharan Africa were established and have captured large amounts of data [33, 34]. There are two main susceptibility bioassays (SB) used for pyrethroid resistance surveillance: the World Health Organisation (WHO) tube protocol [35] and the U.S. Centers for Disease Control and Prevention (CDC) bottle protocol [36]. Under either protocol, offspring of wild mosquitoes are exposed to insecticidal material of a standardised concentration inside a small container for a given amount of time, and the count of dead mosquitoes is recorded. When testing pyrethroid insecticides with the WHO tube protocol, the expose time is 1 hour and a 24 hour post-exposure holding period is applied before mortality assessment. Mosquito samples are obtained by collecting larvae or blood-fed mosquitoes from the field and rearing under standardised conditions. SBs were designed to detect the emergence of resistance in a mosquito population [18, 35] and do not aim to directly capture the impact of resistance on ITN efficacy. The most widely used SB is the discriminating dose bioassay (DD-SB), which operates with a single dose of insecticide to discriminate between fully susceptible and potentially resistant mosquito populations. That discriminating dose, however, does not relate to the insecticide exposure upon interaction with an ITN in the field [13]. Nevertheless, statistical models extrapolating DD-SB data were used to predict the impact of pyrethroid resistance on ITN effectiveness in experimental hut trials [13, 28], and this parameterisation included within transmission dynamics models was broadly able to predict the results of cluster randomised control trials of the mass deployment of different ITNs [37].

To date, resistance to pyrethroids in a mosquito population is typically characterised by a single value, the fraction of mosquitoes dying, and the heterogeneity of tolerance in the population is largely overlooked [9]. In order to better quantify pyrethroid resistance in a mosquito population, intensity-dose bioassays (ID-SB), which assess mosquito susceptibility for a range of insecticide doses, were developed [35, 38] and are increasingly used [39]. A first framework for their analysis proposed fitting a generalised logistic function to ID-SB data and deriving the distribution of lethal doses in the population, from which the lethal dose with respect to any mortality level can be determined [40]. However, these models are non-mechanistic, potentially limiting their practical use for characterising the impact of pyrethroid resistance on disease transmission. In addition, currently no method is available to integrate data from DD-SB and ID-SB to predict the effectiveness of ITNs.

We propose a simple, mechanistic model of insecticide-induced mortality in mosquito bioassays that incorporates stochasticity and explicitly separates the variability in insecticide exposure and in insecticide tolerance. Our model allows us to characterise ID-SB data by two parameters: the median lethal dose (LD_50_), at which half of mosquitoes are killed, and the standard deviation of the logarithm of the lethal dose as a measure of the heterogeneity in insecticide tolerance. While purely statistical models of ID-SB data impose essentially arbitrary functional relationships, the proposed model requires only distributional assumptions on insecticide exposure and tolerance. More generally, it provides a unified treatment of SBs, which are the standard for pyrethroid resistance monitoring, and EHTs, which present a more realistic model system of how mosquitoes interact with insecticides in the field. By fitting our model to the matched SB and EHT data, we obtain a predictive model of the effectiveness of ITN given the SB data.

## Materials and methods

### Data

Following our objective of predicting EHT outcomes from SB data, we only use data from EHT studies where the level of insecticide resistance in the local mosquito population had been characterised in SBs. Both the SB and EHT studies report counts of the number of female *Anopheles* mosquitoes dying, see Fig 1 for a graphical summary. We extended an existing data set of matched DD-SB and EHT studies [28] with a recent study from Burkina Faso comprising EHT and ID-SB data. We included only SB studies on wild-type mosquitoes exposed to single pyrethroid insecticides. All SB studies followed the WHO tube protocol [35], except for one which used a comparable protocol conducted before 2000. We considered only EHT data on single active ingredient, pyrethroid ITNs that were new and unwashed. ITN products which use the same insecticide, but which may differ in other characteristics, were treated the same in our analysis. For the EHT data, we pooled the number of mosquitoes collected across all hut compartments considered in the corresponding study, and mortality counts were determined after a 24-hour recovery period. We did not consider the hut or sleeper volunteer identity as covariates given that previous work indicated that these variables only account for a minor portion of the variation in mosquito mortality across observations [41]. Stratification of mosquito counts by species was not consistent between studies, so for both SB and EHT data, we pooled data for all reported *Anopheles* species. An SB and an EHT study were considered a match if they were conducted in the same village-sized geographical location and in the same year. Within each match, the data were further grouped by insecticide if the insecticides present as active ingredients of the ITNs trialled in the EHT study were all tested in the SB study. If for none of the ITNs the active ingredient was tested in the matched SB study, the data on all the pyrethroid insecticides tested in the SB were pooled, i.e. studies that used different types of pyrethroid were still matched, making the assumption that there is cross resistance within the pyrethroid class [42]. This procedure led to a total of 35 assay pairs of matched SB and EHT data. Control SBs (without insecticide) and EHT control arms (using untreated nets) were included and shared among assay pairs from the same study, without replicating any data. The characteristics and sample sizes per assay pair are listed in Table 1, including references to the original studies. Overall, the data set we used to fit the model includes data for 18,623 mosquitoes observed in SB studies and 110,054 mosquitoes observed in EHT studies (see Table 1 and S6 Table for a summary of all data used).

**Fig 1.**
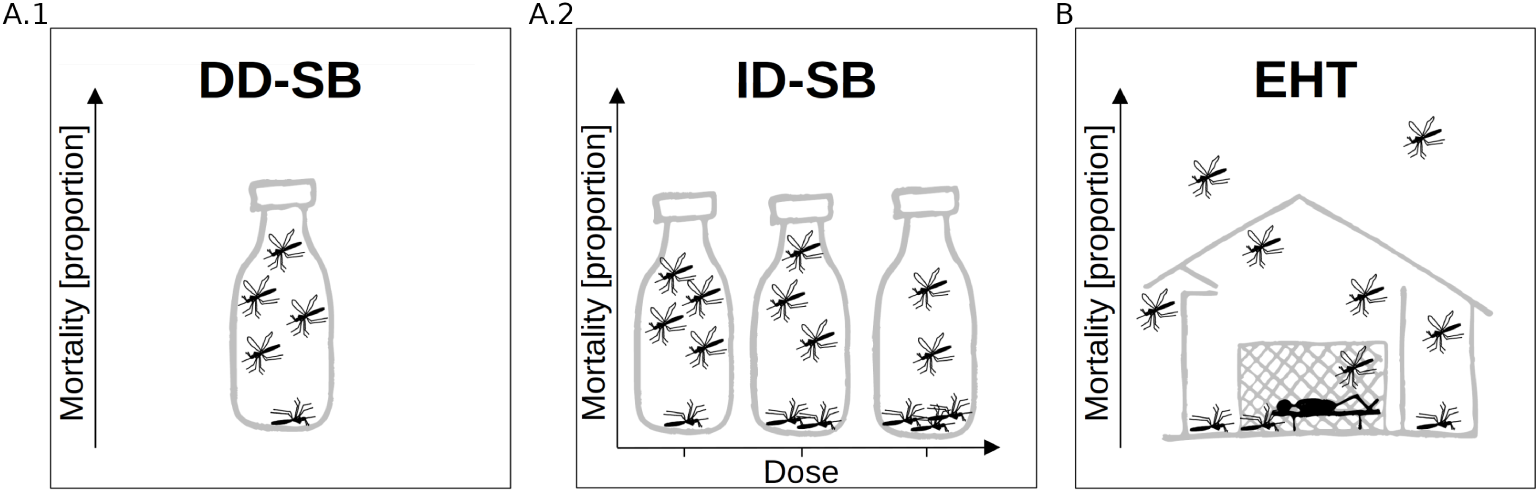
Insecticide bioassay types. **A.1** Discriminating-dose susceptibility bioassay (DD-SB) and **A.2** intensity-dose susceptibility bioassay (ID-SB), both depicted by the CDC bottle bioassay. **B** Experimental hut trial (EHT), depicted by the East African hut type.

**Table 1.** Summary of paired ID-SB and EHT data from Burkina Faso used in this study.

| Assay pair | Site | Year | Insecticide | ITN product | Total in treatment SB | Total in control SB | SB months | Total in intervention hut | Total in control hut | EHT months |
| --- | --- | --- | --- | --- | --- | --- | --- | --- | --- | --- |
| 1 | Tengrela | 2020 | alphacypermethrin | Interceptor | 1214 | 997 | 5-11 | 283 | 731 | 8-9 |
| 2 | Tengrela | 2021 | alphacypermethrin | Interceptor | 804 | 612 | 6-9 | 802 | 886 | 8-9 |
| 3 | Tiefora | 2020 | alphacypermethrin | Interceptor | 1721 | 1376 | 5-11 | 228 | 627 | 8-10 |
| 4 | Tiefora | 2021 | alphacypermethrin | Interceptor | 960 | 929 | 6-9 | 213 | 321 | 8-10 |
| 5 | Tengrela | 2019 | deltamethrin | PermaNet2 | 387 | 318 | 9-11 | 546 | 830 | 9-10 |
| 6 | Tengrela | 2020 | deltamethrin | PermaNet2 | 1353 | 997 | 5-11 | 774 | 731 | 8-11 |
| 7 | Tiefora | 2020 | deltamethrin | PermaNet2 | 1673 | 1376 | 5-11 | 452 | 627 | 8-11 |

All data are from Burkina Faso [43]. Intensity-dose susceptibility bioassays (ID-SB) were conducted according to the WHO tube protocol [35] and experimental hut trials (EHT) used the West African experimental hut design [30]. Note that controls may be shared across assay pairs. For a summary of the paired discriminating-dose susceptibility bioassay (DD-SB) and EHT data additionally used in this study see S6 Table.

### Mechanistic model for insecticide-induced mosquito mortality

Previous models exploring the relationship between insecticide-induced mosquito mortality in the different assays have assumed that all mosquitoes in a DD-SB have the same probability of dying [13, 28]. For ID-SBs, previous work has allowed for heterogeneity in the lethal dose within a mosquito sample, but assumed a uniform exposure to the insecticide [40]. Here we propose a mechanistic model which allows both the insecticide exposure and the maximal insecticide tolerance (measured on ‘Dose’ scale and referred to as the ‘Lethal dose’) to vary between individual mosquitoes by treating those quantities as random variables. Given distributional assumptions, the probability that a mosquito dies depends on assay-specific parameters of the exposure distribution and on the parameters of the lethal dose distribution, which collectively characterise the level of resistance of the corresponding entomological setting. Empirical evidence supports the assumption of heterogeneity in both insecticide exposure and tolerance. For example, variability in insecticide exposure has been observed in confined spaces using video cone tests [44] or in EHT setups using room-scale video tracking [45]. Conversely, a wide section of genetic markers have been associated with pyrethroid-resistant phenotypes in wild mosquitoes [46], and polygenic resistance phenotypes [47] likely exhibit variability in lethal dose. In the absence of data to estimate the correlation between exposure and lethal dose in individual mosquitoes, we make the parsimonious assumption that exposure and lethal dose are independent of each other, while the model would allow for inclusion of a correlation coefficient. The heterogeneity of the exposure varies between bioassays and EHTs, and we assume that the lethal dose distribution for a given mosquito population is the same in both assays. A summary of all model parameters is given in Table 2.

**Table 2.** Summary of model parameters and prior distributions.

| Parameter | Description | Prior distribution |
| --- | --- | --- |
| $\mu_{Tj}$ | mean of log lethal dose in setting $j$ | normal(5, 10) |
| $\sigma_{Tj}$ | SD of log lethal dose in setting $j$ | half-normal(0, 5) |
| $\sigma_{SB}$ | SD of log insecticide exposure in SB | half-normal(0, 5) |
| $\mu_{EHT}$ | mean of log insecticide exposure in EHT | normal(0, 5) |
| $\sigma_{EHT}$ | SD of log insecticide exposure in EHT | half-normal(0, 5) |
| $p^{(0)}_{SB}$ | natural death probability in SB | beta(1, 10) |
| $p^{(0)}_{EHTj}$ | natural death probability in EHT for setting $j$ | beta(1, 10) |

Note that all parameters had independent prior distributions. SD means ‘Standard deviation’.

### Mortality in susceptibility bioassays

Mosquitoes in a SB are heterogeneously exposed to the insecticide due to variation in individual behaviour and we model the exposure heterogeneity stochastically. Let the insecticide exposure of an individual mosquito *i* in a SB be a positive random variable, *X_i_*, with a measuring unit ‘insecticide mass’. We assume *X_i_* = *V_i_c*, where *V_i_* is a positive random variable representing the volume of insecticidal material mosquito *i* comes into contact with and *c* is the average insecticide concentration in a unit volume. Note that we use the term volume in a broad sense, including all uptake of insecticidal material during the bioassay, such as through surface contact with the insecticide carrier and through contact with vaporised insecticide in the air. We further assume that the concentration *c* is a linear function *c* = *αd* of the dose *d* of insecticide applied to the carrier in the SB, with the dose *d* measured as multiples of the discriminating dose defined by the WHO [35]. We model *V_i_* as log-normally distributed to account for the likely large variability in mosquito exposure to insecticide in the SB. By appropriately rescaling the volume measuring unit – corresponding to e.g. switching from inches^3^ to centimeters^3^ – we can assume log(*V_i_*) has mean − log(*α*) without loss of generality. Hence, the exposure *X_i_* = *αdV_i_* follows a log-normal distribution and due to the rescaling *α* vanishes from the equation:

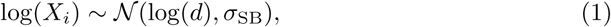

where *σ*_SB_ *>* 0. The measurement unit for exposure is ‘insecticide mass’ but due to the change in the volume scale, it has the same scale as ‘Dose’, measured as multiples of the discriminating dose defined by the WHO [35].

We represent the lethal dose for mosquito *i* as a positive random variable *T_i_* and assume that mosquito *i* dies in the 24 hours post exposure (the holding time of mosquitoes in both SB and EHT for insecticides with a fast mode of action, such as the pyrethroid products considered here) if *X_i_ > T_i_*. The indicator random variable for death due to the insecticide **1***_Xi>Ti_* has a Bernoulli distribution with probability

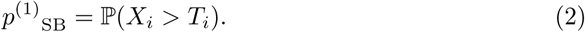

However, mosquitoes can also die due to other exogenous factors during the holding time of the assay, and we assume an additional independent Bernoulli death process with probability *p*^(0)^_SB_. By the law of total probability, the death of mosquito *i* is a Bernoulli random variable with probability

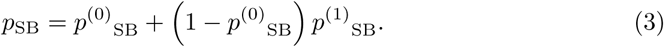

### Mortality in susceptibility bioassays for field-collected mosquitoes

Mosquitoes sampled from a wild population exhibit variable insecticide susceptibility, due to genetic variation and variation in environmental conditions during both juvenile and adult life stages. We assume that the lethal dose *T_i_* for an individual mosquito *i* is independent of its exposure *X_i_* and is distributed following a log-normal distribution

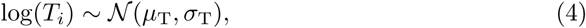

where the mean *µ*_T_ and the standard deviation *σ*_T_ characterise the lethal dose distribution within a given population. By the monotonicity of the log function and the symmetry of the standard normal distribution, the probability of insecticide-induced death can be computed as

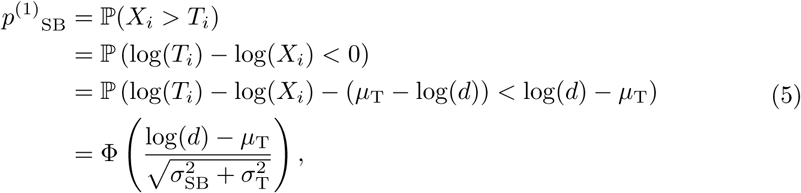

where Φ denotes the cumulative distribution function of the standard normal. To derive Eq. 5, we used the fact that the sum of two independent normal distributed random variables is normally distributed with mean equal to the sum of the two means and variance equal to the sum of the variances. For an illustration of Eq. 5 see Fig 2.

**Fig 2.**
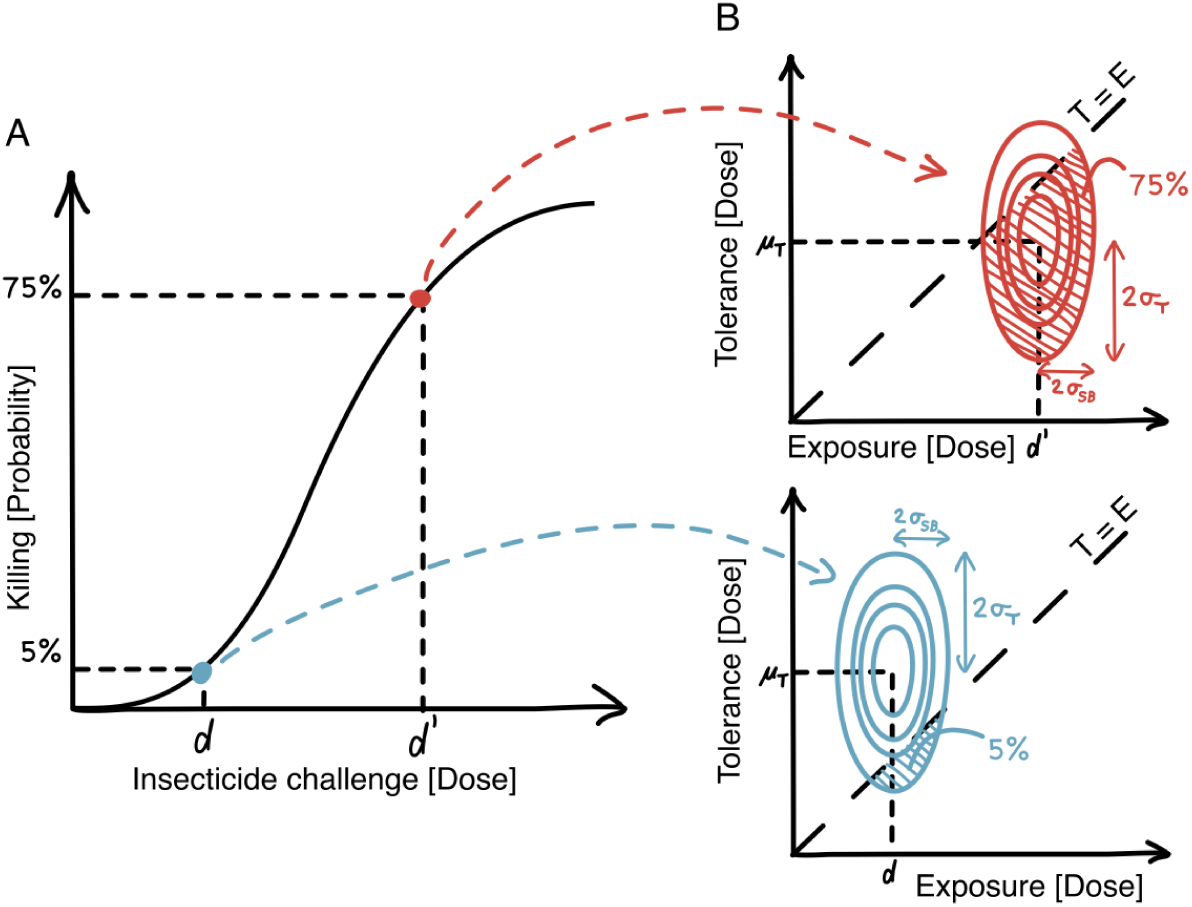
Our mechanistic model for insecticide-induced mosquito mortality in a ID-SB. **A** Illustration of the modelled relation between the insecticide dose tested in a ID-SB (‘insecticide challenge’) and the probability for killing a mosquito from a given population. **B** Illustration of how the killing probability is computed for a low dose *d* (blue) and high dose *d*^′^ (red) used to challenge mosquitoes in a ID-SBs. The parameters *µ*_T_ and *σ*_T_ define the log-normal distribution of the lethal dose in the mosquito population, i.e. the maximum tolerance, and the challenged dose together with the parameter *σ*_SB_ define the log-normal distribution of the insecticide exposure in the SB. The joint distribution of exposure and lethal dose take the form of a bivariate normal when plotted on the logarithmic scale, here displayed as contour lines. The killing probability for a given insecticide challenge is given by the probability that the exposure exceeds the lethal dose, i.e. the maximum tolerance (*E > T*).

From Eq. 5 it follows that LD_50_, the dose at which 50% of the mosquitoes die, is given by exp(*µ*_T_) in our model, the median of *T_i_*. This is due to the symmetry of the exposure and lethal dose distributions on the logarithmic scale and the monotonicity of the logarithm. The parameters *σ*_T_ and *σ*_SB_ define the shape of the dose-response curve; see Fig 3 for a an illustration. This model allows us to characterise the insecticide resistance of a mosquito population by location and scale parameters: the median lethal dose (exp(*µ*_T_)), and the standard deviation of the logarithm of the lethal dose (*σ*_T_) as a measure of the heterogeneity in insecticide tolerance.

**Fig 3.**
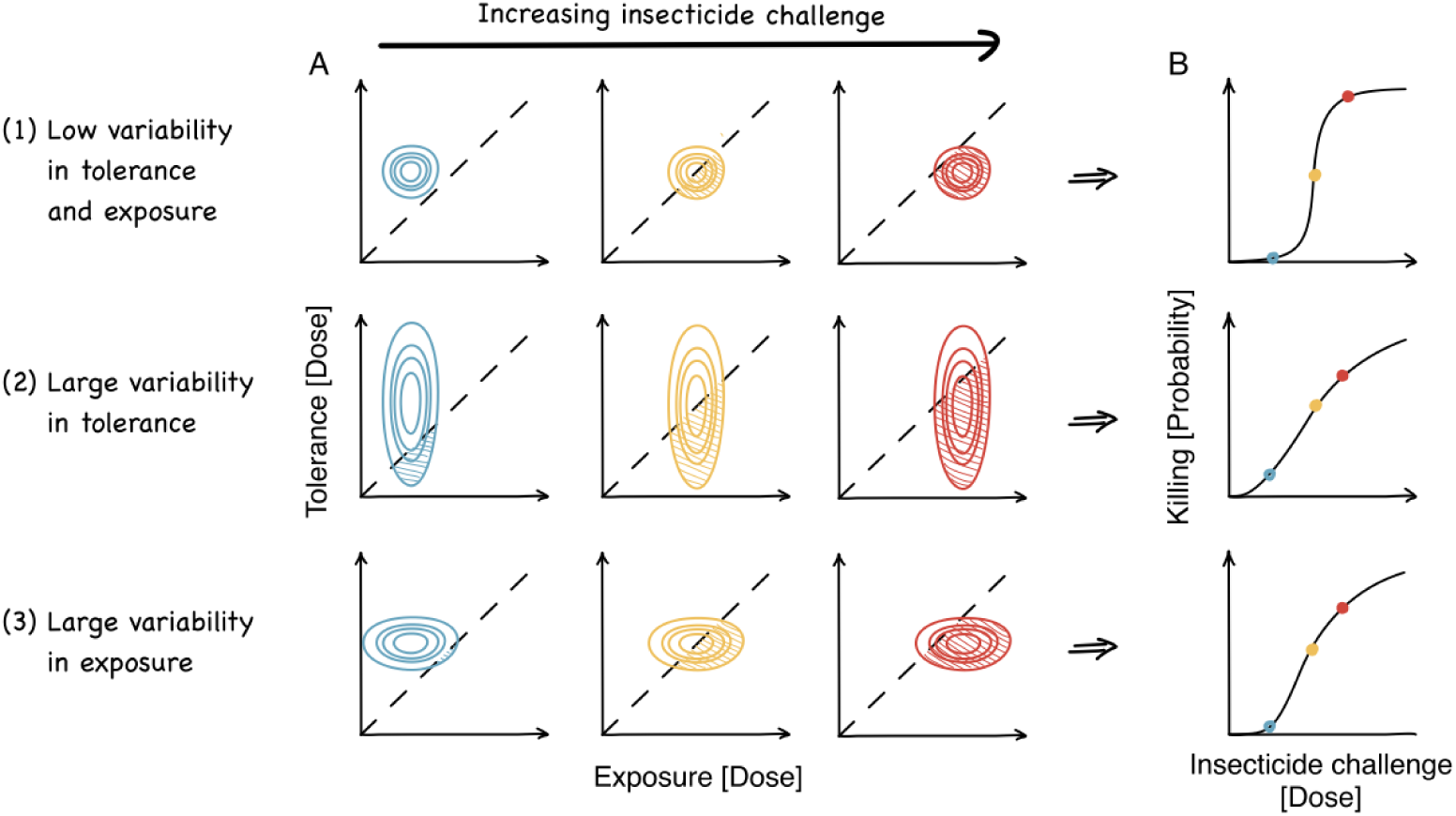
How variation in insecticide tolerance and exposure impact the dose-response relation under our mechanistic model. For the three scenarios of (1) low variability in both tolerance and exposure, (2) large variability in tolerance, and (3) large variability in exposure, we display **A** the bivariate distribution of exposure and lethal dose, i.e. the maximal tolerance, on the logarithmic scale, for three levels of insecticide challenge, and **B** the corresponding dose-response curve of the mosquito killing probability for increasing insecticide challenge on the actual scale.

### Mortality in experimental hut trials

For EHTs we use a similar model. We assume that mosquitoes from the local population enter the experimental hut randomly and independently of their individual lethal dose. Note that this means that we assume that there is no behavioural resistance to the insecticide, and that each mosquito entering the hut has a lethal dose *T_i_* distributed as in Eq. 4 with parameters *µ*_T_ and *σ*_T_ corresponding to the local population. We then assume that each mosquito *i* in the hut experiences an exposure *X_i_* that is independent of its lethal dose *T_i_* and distributed according to

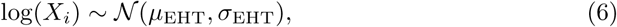

with parameters *µ*_EHT_ and *σ*_EHT_ *>* 0. Like in SBs, we assume that a mosquito *i* dies if *X_i_ > T_i_*and by Eq. 2 the probability of death due to insecticide is

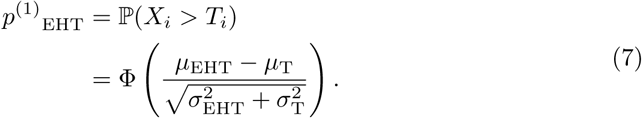

Similarly to Eq. 3, we assume an independent background mortality process with probability *p*^(0)^_EHT_ of death for a given EHT study. Again, the death of mosquito *i* is a Bernoulli random variable with probability

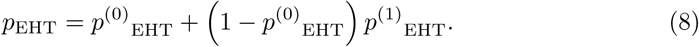

### Modelling the counts of dead mosquitoes in susceptibility bioassays and experimental hut trials

Assuming insecticide exposure and lethal dose are conditionally independent given the bioassay type and the resistance setting, respectively, the count of mosquitoes dying in a replicate of the assay follows a binomial distribution. So for each assay pair, *j*, corresponding to all SB and EHT data from a given village-sized geographical location and a given year (for details on matching SB and EHT studies see the Data section), we can derive the following sampling distributions. For the SBs, we have

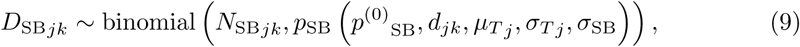

where *D*_SB*jk*_ is the count of dead mosquitoes and *N*_SB*jk*_ the total number of mosquitoes tested in the *k*th SB replicate of assay pair *j*. The probability of death *p*_SB_ in the corresponding SB replicate is a function of the insecticide dose *d_jk_* applied in the SB replicate and the parameters described in Table 2, as defined in Eqs 3 and 5. For the corresponding EHT, we have

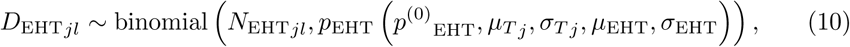

where *D*_EHT*jl*_ is the count of dead mosquitoes and *N*_EHT*jl*_ the total number of mosquito captured in the hut in the *l*th EHT replicate of assay pair *j*. The death probability *p*_EHT_ in the corresponding EHT replicate is a function of the parameters described in Table 2, as defined by Eqs 7 and 8.

Taken together, the models defined in Eqs 9 and 10 form the joint model across different bioassays for a given resistance setting. The two sub-models are coupled via the common parameters *µ*_T_*_j_* and *σ*_T*j*_ per assay pair *j*, which completely define the lethal dose distribution for the corresponding resistance setting. The exposure parameters *σ*_SB_, *µ*_EHT_, and *σ*_EHT_ are fixed across all assay pairs (see the discussion for associated model limitations). By coupling of the two sub-models (Eqs. 9 and 10), the exposure in EHTs is defined on the same scale – ‘Dose’ – as the exposure in the SBs. Although we estimated independent natural death probabilities in the EHTs for each assay pair *j* (*p*^(0)^_EHT_*_j_*), we assumed the same natural death probability in the SBs across all assay pairs (*p*^(0)^_SB_) since for the DD-SB studies there were no control data available. Assuming conditional independence across all assay pairs, we multiply the probabilities corresponding to Eqs. 9 and 10 to produce the sampling distribution for all data in Table 1 and S6 Table.

### Predicting the killing effect of ITNs from intensity dose susceptibility bioassays

The objective of this study is to provide a model for predicting the effectiveness of an ITN in killing mosquitoes in a given setting, as measured in an EHT, directly from ID-SBs with the corresponding insecticide in the same setting. After calibrating the parameters *σ*_SB_, *µ*_EHT_ and *σ*_EHT_ of the joint model with the present data, the resistance characteristics (*µ*_T∗_*, σ*_T∗_) of a previously unknown setting ∗ may be estimated from new ID-SB data by the SB sub-model (Eq. 9). These estimates may then be plugged into the EHT sub-model (Eq. 10) to predict the killing effect of an ITN with the corresponding insecticide in a hypothetical EHT, as illustrated in S1 Fig for an example setting. We assessed the predictive accuracy of this approach using leave-one-assay-pair-out cross-validation. Specifically, we left out, one by one, the EHT intervention arm data points for each of the assay pairs with ID-SB data (see Table 1), fitted the joint model to the rest of the data (including the DD-SB assay pairs) and then predicted the left-out EHT mortality probabilities from the estimated IR metric, EHT exposure and replicate-specific EHT control mortalities. This procedure allows for a fully Bayesian prediction, taking advantage of the fact that the posterior samples for the tolerance and exposure parameters are naturally matched by Stan iteration in the joint model.

### Bayesian inference

The joint model composed of the two sub-models (Eqs. 9 and 10) was fitted to the full data set (Table 1). In addition, the SB model (Eqs. 9) was fitted separately to the ID-SB data (Table 1) with independent natural death probabilities *p*^(0)^_SB_*_j_* per assay pair *j*. We fitted the models using a Bayesian framework, with prior distributions shown in Table 2. These were generally weakly informative (S2 Fig shows prior predictive checks) with the exception of the prior for the daily background mortality probability (i.e., mosquito mortality in the presence of no insecticide) which we assumed had 65% prior probability mass below 10% mortality.

Both models were fitted using the probabilistic language Stan [48] and the posteriors were sampled using Stan’s default NUTS algorithm [49] accessed through the rstan package v2.26.1 [50] in R version 4.4.0 [51]. For the SB model and the joint model, we ran 4 chains with 4, 000 and 6, 000 iterations, respectively, and each time discarded half of the iterations as warm-up. To reduce the need for warm-up, we initialised the chains with point estimates obtained by optimisation; to obtain these point estimates, we used Stan’s L-BFGS algorithm and used the set with the highest log-posterior across 10 optimisation runs. For processing the Stan output, we used the tidybayes R package [52]. For all model fits, we diagnosed convergence with *R̂* < 1.01 and bulk- and tail-effective sample sizes above 400 for all model parameters [53]. For making statements about estimated quantities with a certain credibility, we relied on central credible intervals (i.e. equally tailed quantile intervals) for univariate quantities because of their invariance with respect to monotonic transformations, and on highest density credible regions for multivariate quantities because of the lack of a natural choice for the quantile function for a multi-variate random variable.

### Reproducibility

The data and code used to generate the presented results are publicly available [54]. The code is released under the MIT licence. Relevant software and packages are mentioned in the corresponding methods sections. All analyses were conducted in a Linux environment (Ubuntu 22.04.4). Inference with Hamiltonian Monte Carlo methods is inherently stochastic, and, while we fixed and reported the seeds for the random number generation, running the same code on another computing environment may lead to minorly different parameter estimates.

## Results

### The mechanistic model can integrate results from different bioassays

Fig 4 shows that the joint model (Eqs 9 and 10) fit simultaneously to all SB and EHT data recovered the mortality trends in both assays. Numerical summaries of the posterior predictive ITN killing effect in EHTs are contained in S9 Table. The model-based estimates are reasonable even for the studies with single-dose DD-SB for which the two-parameter resistance characterisation typically is not well identifiable (indicated by large posterior parameter correlation; see S9 Table). The estimates are generally a better fit for SB than for EHT studies, which was to be expected since, in the former experiment, conditions are more controlled. When the model-based estimates of EHT mortality are far from the observed mortality probability, the uncertainty of the latter is generally large (Fig 4). Inferred insecticide resistance characteristics and ITN killing effects for all assay pairs are shown in S3 Fig and summarised numerically in S9 Table; the inference for all parameters of the underlying model is summarised in S7 Table. For the predictive accuracy of mortality predictions for the EHT data from Burkina Faso using ID-SB data only (leave-one-assay-pair-out cross-validation), see S4 Fig corresponding to Fig 4B.

**Fig 4.**
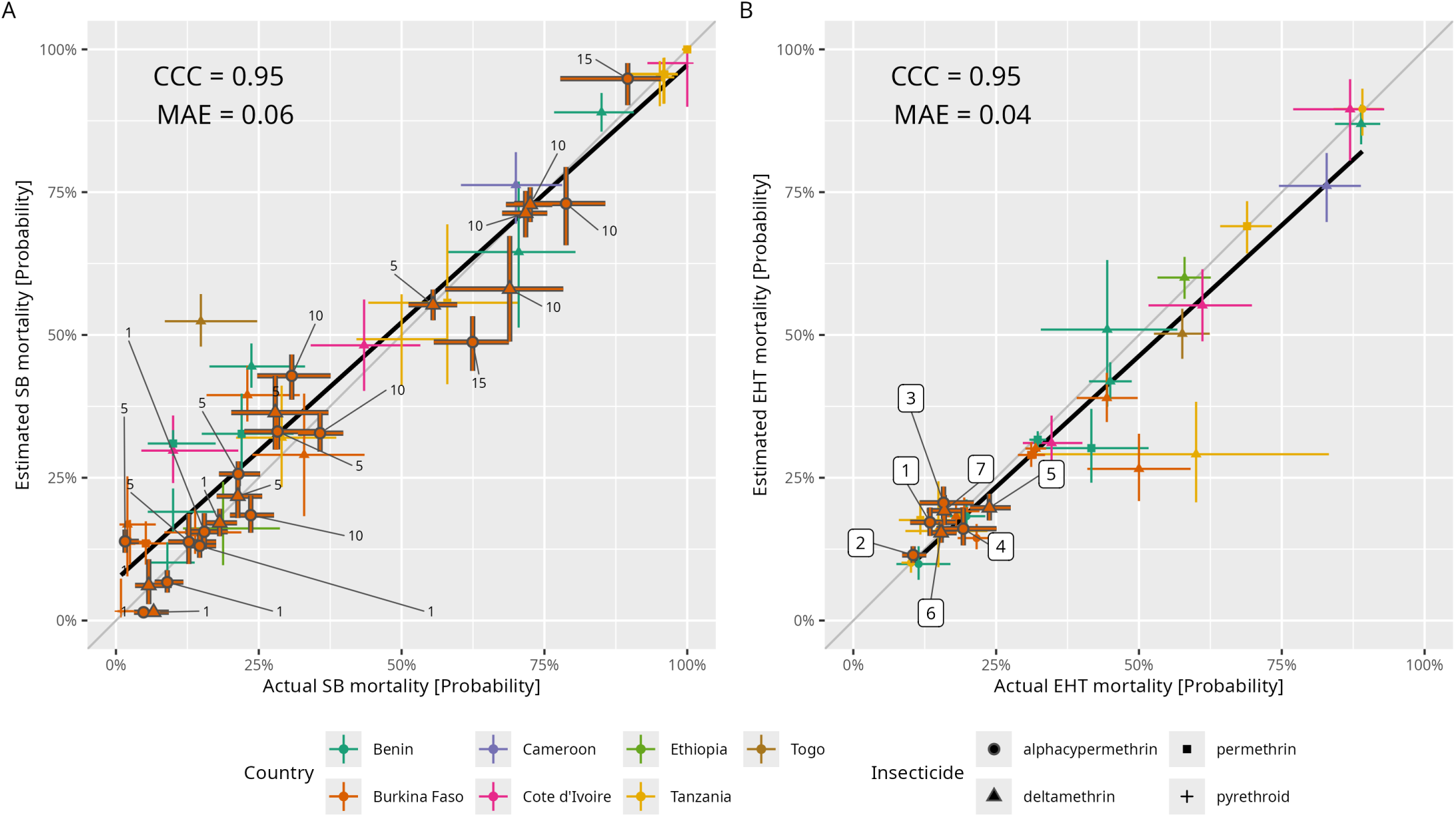
Model estimates of mosquito mortality in susceptibility bioassays and experimental hut trials. Mortality in **A** susceptibility bioassays (SBs) and **B** experimental hut trials (EHTs), for 35 different assay pairs matched by geographical location, year and insecticide. Assay pairs comprising intensity-dose SBs (ID-SBs) are outlined in dark grey, with numeric labels in panel A indicating the dose and numeric labels in panel B indicating the assay pair number (see Table 1). For each assay pair, and each dose for ID-SBs in panel A, the mortality probability estimated by a binomial model with a uniform prior probability (‘actual’) is plotted against the mortality probability estimated by the joint mechanistic model fitted to all SB and EHT data jointly (‘estimated’). See S4 Fig for predictions of EHT mortality from ID-SB data only using the joint model calibrated to all SB and EHT data except the EHT intervention arm of the given assay pair (leave-one-assay-pair-out cross-validation). Point estimates correspond to the posterior median and interval estimates to 95% central, quantile-based credible intervals. A linear regression of the posterior medians (point estimates, neglecting uncertainty) is plotted in black to visualise the actual versus estimated fit. Correspondingly, Lin’s concordance correlation coefficient (CCC) measures agreement of the point estimates with the identity line on a scale between -1 and 1 (with 1 indicating perfect alignment), and the mean absolute error (MAE) indicates the average absolute error of the model-estimated mortality on the probability scale. The insecticide was recorded as ‘pyrethroid’ if different insecticides were pooled for the corresponding assay pair (see Data section and Table 1).

### Varied insecticide susceptibility dose-response patterns are captured by the mechanistic model

Our mechanistic model could represent the ID-SB data patterns from Burkina Faso, which comprised two locations over three years and two insecticides, as shown by the estimated dose-response curves in Fig 5. However, for two resistance settings (assay pairs 1 and 4), the model estimates did not align well with the data since our mechanistic model only supports S-shaped dose-response curves. In addition to the joint model fit to all data, we fitted the SB model (Eq. 9) to the ID-SB data only (see S8 Table for parameter inference). Comparison of the dose-response curves (Fig 5) and the underlying resistance metrics (S7 Table and S8 Table) across the two fits suggests that the resistance metric is inferred from the ID-SB data with limited influence from the EHT data in the joint model fit. This is important since, in the actual use case, we want to use the joint model to predict the ITN killing probability in a hypothetical EHT from ID-SB data only (see also the leave-one-assay-pair-out cross-validation, S4 Fig).

**Fig 5.**
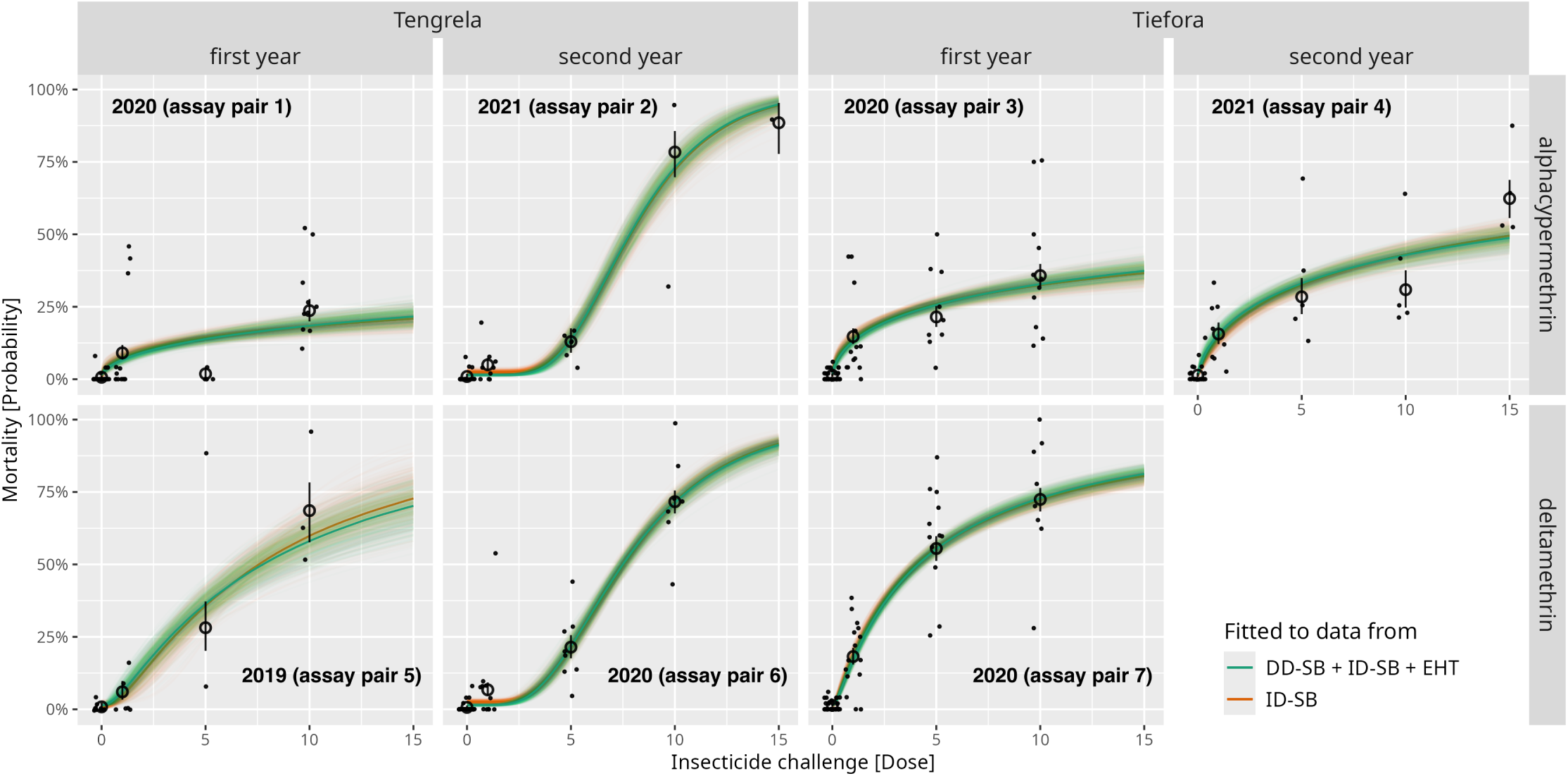
Model fits to intensity-dose susceptibility bioassay data from Burkina Faso. Each panel corresponds to one of the assay pairs which included intensity-dose susceptibility bioassay (ID-SB) data (see Table 1). Each panel reports the observed mortality percentages in the corresponding ID-SB replicates across the tested doses (black points, plotted with random jitter), the median mortality per dose (black circles), and the 95% central credible intervals of the mortality probability per dose as estimated with a separate binomial model per assay pair and dose (black vertical bars). ‘Dose’ refers to multiples of the discriminating dose defined by the WHO [35]. Note that for deltamethrin, doses 1, 5 and 10 were tested, and for alphacypermethrin there was an additional dose at 15. The underlaid dose-response curves correspond to the estimates of the mechanistic model for mortality in ID-SBs, either fitted simultaneously together with the experimental hut trial (EHT) model to all DD-SB and EHT data (green) or fitted separately only to ID-SB data (orange). The thick dose-response curves are the median posterior predictive model estimates with respect to the two fits; corresponding uncertainty is depicted by 500 curves that were generated with parameter sets sampled randomly from the corresponding posteriors. Summaries of the posteriors for all parameters underlying the two model estimates are contained in S7 Table and S8 Table.

### Quantifying differences in insecticide exposure across assays

Our mechanistic model allows for a comparison of insecticide exposure in SBs compared to EHTs. The estimated exposure distributions in Fig 6A indicate that exposure in EHTs was markedly lower and slightly more variable than in SBs. In particular, the median exposure in EHTs was 0.50 [0.39, 0.62] (posterior median and 95% credible interval) times the discriminating dose defined by the WHO, and the standard deviation of the exposure measured on the logarithmic scale was 0.15 [0.02, 0.29] in EHTs compared to 0.11 [0.01, 0.24] in SBs.

**Fig 6.**
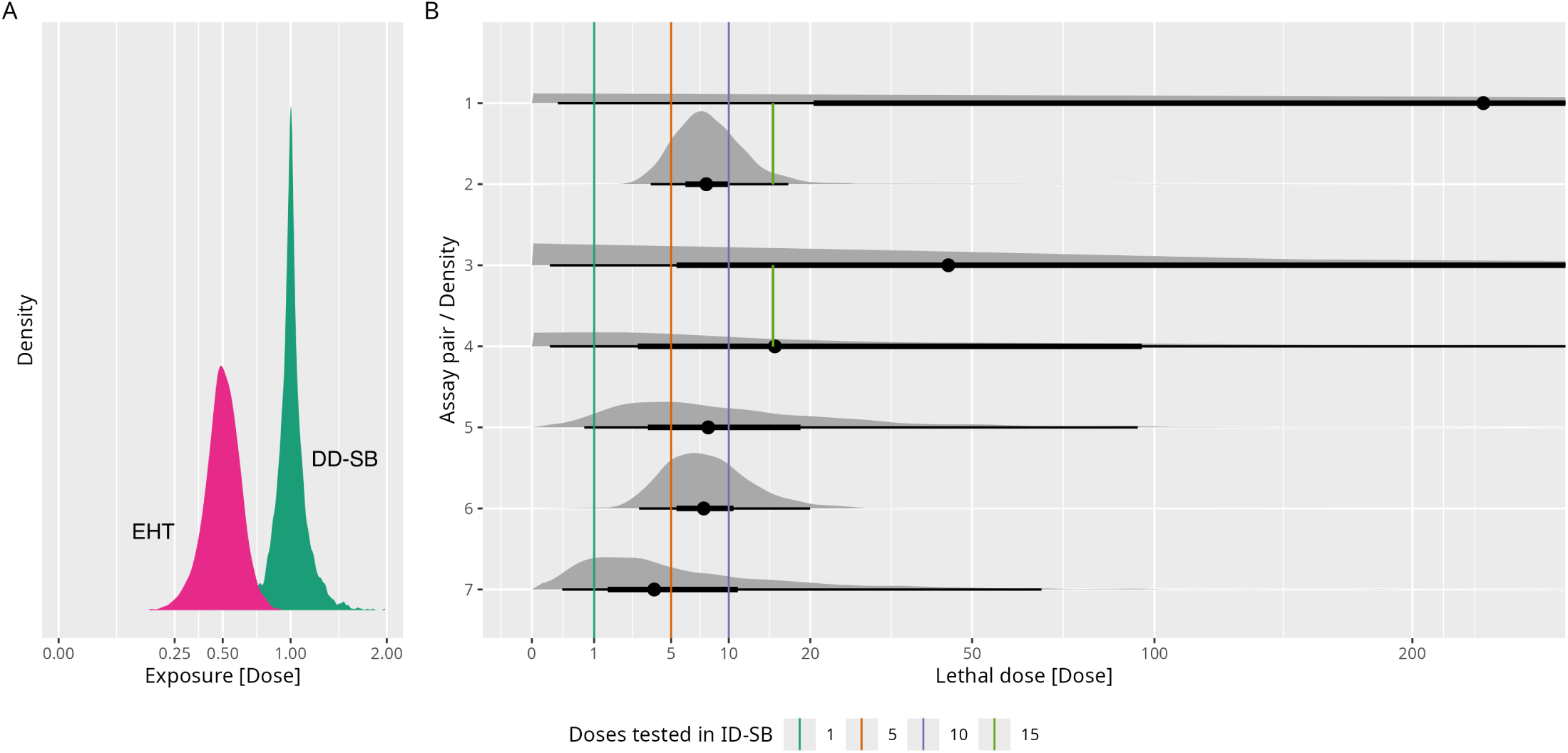
Model estimates of insecticide exposure in susceptibility bioassays and experimental hut trials, and minimal lethal dose across resistance settings. **A**Posterior predictive distributions of exposure to pyrethroid insecticides in susceptibility bioassays (SBs) and experimental hut trials (EHTs) on the ‘Dose’ scale (multiples of the discriminating dose defined by the WHO [35]), as estimated by the joint mechanistic model. We assumed the same variance of the log-exposure distribution across discriminating-dose SBs (DD-SBs) and intensity-dose SBs (ID-SBs), so that the depicted DD-SB exposure distribution (green) represents the exposure variability across all considered SBs. For ID-SBs, the log-exposure distribution is shifted to be centred at the corresponding dose, with equal variance. The horizontal axis is on a square root scale. **B** Posterior predictive lethal dose distributions across all resistance settings with ID-SB data (assay pairs 1-7, see Tab. 1). Grey areas depict densities, black points denote medians and black lines show central, quantile-based credible intervals for probability levels of 50% and 95%. Vertical lines indicate the doses – which equal the median exposure in the ID-SB – against which the lethal dose distribution of the population was tested in the ID-SBs. The horizontal axis is on a square root scale. To facilitate the density estimation for this plot, we removed the top 0.05% percentile of the sampling points.

### Improved characterisation of resistance using estimates of tolerance heterogeneity

We characterise the resistance of a local mosquito population against a given insecticide by the heterogeneity of insecticide tolerance, i.e. the lethal dose distribution in the population, in addition to the median lethal dose. This two-parameter resistance metric allows for a more detailed comparison of resistance profiles across different locations, years, and pyrethroid types. Fig 7A graphically displays the estimated resistance characteristics for the assay pairs comprising ID-SBs, including uncertainty quantification (highest density credible regions). The two parameters could be inferred from ID-SB data, but estimates were highly correlated if the insecticide challenge did not explore the full mortality range (assay pairs 1, 3 and 4, see S9 Table). Fig 6B shows the corresponding lethal dose distributions, which are markedly more dispersed for assay pairs 1, 3, and 4 compared to the others, due to the higher heterogeneity in insecticide tolerance and the more uncertain median lethal dose estimates. For an improved estimation of the two-parameter resistance metric from future ID-SB studies, we recommend using dose escalation up to about 80% mortality (LD80), including a control and the discriminating dose, as suggested by a simulation study informed by the present ID-SB data from Burkina Faso (see S5 Fig). The resistance characteristics for the assay pairs that comprise only DD-SBs are depicted in S3 Fig A and summarised numerically in S9 Table, but these estimates were typically highly correlated and uncertain.

**Fig 7.**
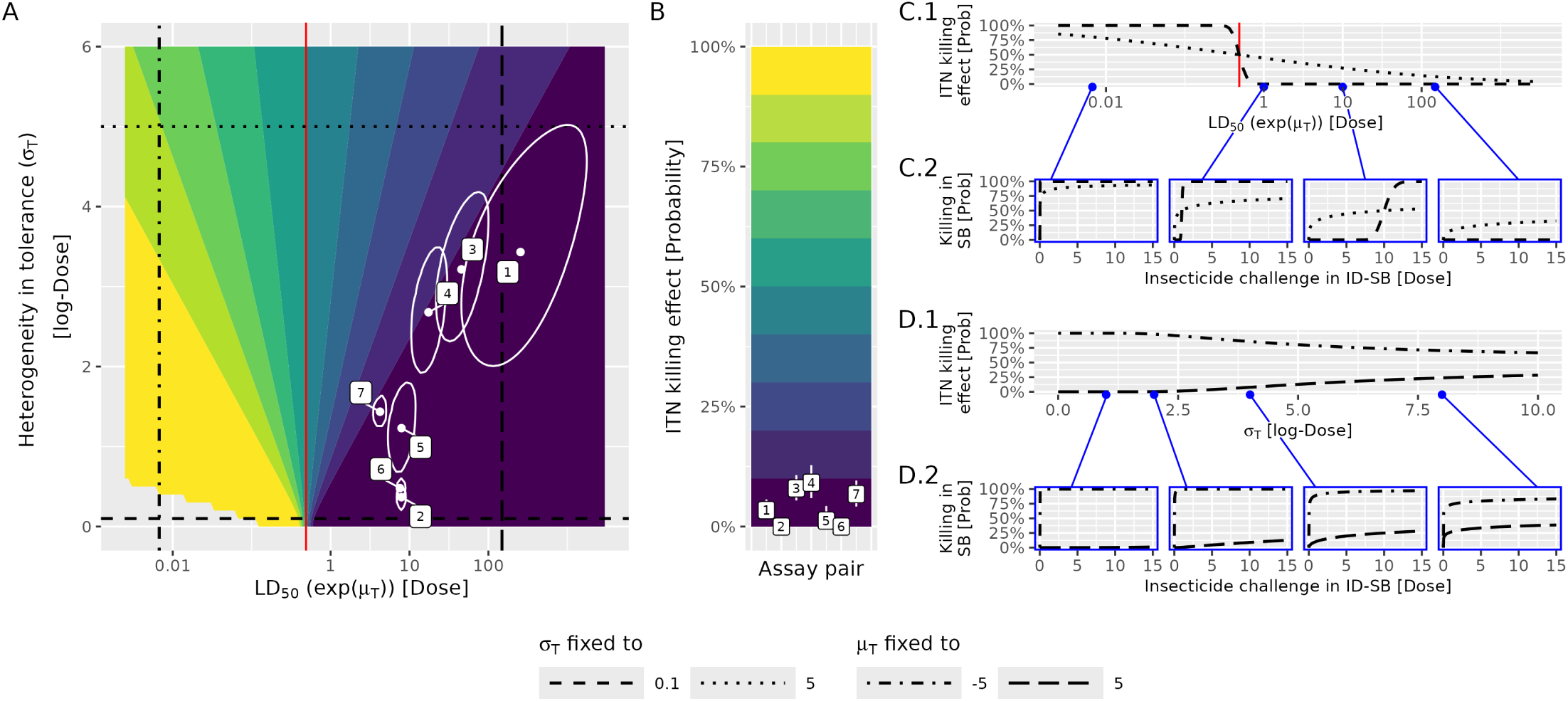
Estimated insecticide resistance metrics and ITN killing effects, and projected ITN killing effects under varying insecticide resistance metrics. **A** Insecticide resistance characterisations by the median lethal dose (LD_50_, equal to exp(*µ*_T_)) and the proposed tolerance heterogeneity measure (standard deviation of the logarithm of the lethal dose, *σ*_T_) for all assay pairs which comprised intensity-dose susceptibility bioassays (ID-SB) (assay pairs 1–7, see Table 1). Labelled dots (white) represent posterior medians, and curves (white) encompass 95% highest density credible regions. Numerical summaries of the estimated resistance characteristics are contained in S9 Table. Coloured areas correspond to binned posterior predictive ITN killing probabilities (posterior median, see panel B for colour legend), as estimated by the joint mechanistic model. The red vertical line marks the estimated median insecticide exposure in an EHT (posterior median). Black horizontal lines represent transects through the parameter space at *σ*_T_ values of 0.1 and 5, and black vertical lines transects at LD_50_ values of 0.007 and 148.4 (*µ*_T_ values of −5 and 5, respectively), with corresponding ITN killing effects and ID-SB dose-response curves displayed in panels C and D. **B** Posterior predictive ITN killing probabilities corresponding to the resistance settings characterised in A (assay pairs 1-7, see Table 1), displayed by the posterior median (square white markers) and central, quantile-based 95% credible intervals (vertical white lines). Numerical summaries of the estimated ITN killing probabilities are contained in S9 Table. **C.1** Posterior predictive ITN killing effect when varying the median lethal dose (LD_50_) while fixing the tolerance heterogeneity (*σ*_T_) to either 0.1 or 5 (dashed and dotted curve, respectively, corresponding to the horizontal transects in panel A). The red vertical line marks the estimated median insecticide exposure in EHT (posterior median). **C.2** Corresponding example dose-response curves for selected LD_50_ values of 0.007, 1, 10 and 148.4 (corresponding to *µ*_T_ values of −5, 0, log(10) and 5, respectively). **D.1** Posterior predictive ITN killing effects when varying the tolerance heterogeneity (*σ*_T_), while fixing the median lethal dose (LD_50_) to either 0.007 (*µ*_T_ = −5) or 148.4 (*µ*_T_ = 5) (dot-dashed and long-dashed curve, respectively, corresponding to the vertical transects in panel A). **D.2** Corresponding example dose-response curves for selected *σ*_T_ values of 1, 2, 4 and 8. ‘Dose’ refers to multiples of the discriminating dose defined by the WHO [35].

Across both locations and both insecticides, we found that the lethal dose was well beyond the insecticide exposure in an EHT for most mosquitoes (compare the lethal dose distributions in Fig 6B with the EHT exposure distribution in Fig 6A), and the ITN killing effects were therefore very low (Fig 7B). The incomplete dose escalation for the assay pairs 1, 3, and 4 made comparison of the tolerance distributions difficult. We could not detect a consistent inter-location difference in resistance characteristics between the two villages, Tengrela and Tiefora, in Burkina Faso. With respect to intra-location resistance dynamics, we estimated that in Tengrela tolerance heterogeneity against both insecticides decreased over the two years (assay pairs 1 versus 2 for alphacypermethrin and 5 versus 6 for deltamethrin, numerical results in S9 Table). We inferred that the lethal dose distribution was considerably wider for alphacypermethrin than for deltamethrin in both locations (assay pairs 1 versus 6 for Tengrela and 3 versus 7 for Tiefora).

### Predicting ITN killing effects across insecticide resistance metrics

In Fig 7 C and D we predict the ITN killing effect for a range of possible resistance characteristics, given the calibrated joint model (median posterior predictive killing probability). As expected, the ITN killing effect decreases with increasing median lethal dose (LD_50_), with a decrease rate depending on the tolerance heterogeneity measure (*σ*_T_) (Fig 7 C.1 and C.2). Regardless of the *σ*_T_ value, a 50% ITN killing effect is achieved if the median lethal dose is the same as the median exposure in the EHT. Varying the tolerance heterogeneity measure (*σ*_T_) shows a more nuanced effect on EHT mortality (Fig 7 D.1 and D.2): Low heterogeneity (small *σ*_T_ values) accentuate extremes in the ITN killing effect for low and high *µ*_T_. High heterogeneity (large *σ*_T_ values) pushes the ITN killing effect towards 50% since it produces highly dispersed lethal dose distributions.

## Discussion

We presented a mechanistic model of insecticide-induced mosquito mortality in bioassays that explicitly describes the interaction of mosquitoes with the insecticide. By fitting the model to data from a range of malarial locations in Sub-Saharan Africa, we have illustrated that the model provides a common framework for interpreting and comparing data from a range of phenotypic insecticide bioassays. Namely, we proposed a two-parameter resistance metric, given by the median lethal dose and a measure of heterogeneity in insecticide tolerance, for improved characterisation of resistance profiles across sites. Generally, the larger the heterogeneity measure, the more robust the ITN killing effect with respect to variation of the insecticide exposure, for example, due to differences in housing structures.

EHTs are thought to provide a realistic emulation of the performance of ITNs in the field, and a direct outcome of them is a measure of the killing effect of ITNs. Our model provides a framework for using data collected from considerably cheaper ID-SBs to emulate the results of a hypothetical EHT. We have tested this approach using cross-validation on a data set from Burkina Faso. However, our estimated ITN killing effects for the assay pairs comprising ID-SB data were consistently low, corresponding to high levels of resistance in the relatively few studies where both ID-SB and EHT data were available. Our model needs to be reassessed once data from a more diverse range of sites become available. This will allow exploration of the levels of exposure observed in EHTs with different structural designs to allow more reliable comparisons of different products and mosquito populations between sites.

Previously, non-mechanistic models have been used to infer a heuristic link between mortality in DD-SBs and EHTs [13, 28], and these estimates were subject to large uncertainty. In response to the need for new resistance testing guidelines to better inform control programmes [55], ID-SBs were developed [39] and the dose-mortality relationship has been analysed using flexible statistical models [40]. Our characterisation of resistance is similar to the lethal concentration estimated in [40], but the approaches differ in the underlying model. In [40], the lethal concentration distribution is defined by fitting a flexible sigmoidal curve (a generalised logistic function of five parameters) to the ID-SB data and interpreting that curve as the cumulative distribution function of the lethal concentration. This means that the shape of the dose-response curve is entirely reflected by the lethal concentration distribution, implicitly assuming identical exposure of all mosquitoes. In contrast, we allow for stochastic exposure and, under some distributional assumptions, we derive a cumulative distribution function for mortality. This means that the shape of our dose-responses curve is described by both the variance of the lethal dose and the exposure in the mosquito population, while the dose where the curve reaches 50% mortality is entirely determined by the median lethal dose. Despite its mathematical simplicity, our model fits the ID-SB data well relative to these more heuristic statistical approaches, and its mechanistic nature provides greater interpretability.

Here, we focus on the killing effect of ITNs as measured in EHTs and make no statements about house entry and feeding inhibition effects of ITNs. When predicting the ITN killing effect, we assume that the IR metric estimated from SB data is independent of house entry rates since the present data do not allow us to estimate a potential relation. This does not mean that we assume there is no behavioural resistance, only that it cannot be detected from SB data. For our killing estimates, we do not explicitly account for any factors beyond our IR metric. Thus, factors that were present during the matched SB study, such as microbial infections in the local mosquito population, are implicitly accounted for when predicting ITN killing effects from SB data. However, factors that varied between the SB and EHT studies of a matched assay pair – such as environmental conditions (including during the juvenile mosquito stages) and very localised conditions at the EHT site, such as sugar feeding opportunities – are not accounted for, potentially impacting predictive performance. Also, we have assumed uniform insecticide exposure across all EHT data, while studies differ in hut design, seasonality and mosquito species composition. The EHT study with concurrent ID-SB data from Burkina Faso used a single hut design and was run during the rainy season, with some variation in study months between SB and EHT as well as between assay pairs (see Table 1). Furthermore, we currently treat data from two different years from the same geographic location as two independent entomological settings. Future work with a larger and richer data set is needed to further investigate the sources of variation in bioassay and EHT results.

Due to data limitations, we have not distinguished between different *Anopheles* species. Our modelling approach does not require all mosquitoes in a sample to be from the same species, but does require that the SB sample represents the EHT sample within a given assay pair. This, as well as our assumption of uniform exposure across EHT studies, likely affected our goodness of fit. Also, the ability of our model to estimate the heterogeneity of insecticide tolerance in a local mosquito population is limited by the efforts made to obtain a representative mosquito sample. Hence, we may underestimate heterogeneity if mosquito samples contain siblings. Both species composition and variation in exposure between EHT studies could be addressed with a hierarchical model extension when a richer data set including ID-SB data becomes available.

For the calibration of the exposure parameters, we fitted a joint model simultaneously to the SB and EHT data. This is problematic since the resistance characterisation parameters are likely informed by both the SB and the EHT data, while they must be inferred from the SB data alone when making predictions of ITN effectiveness for settings without EHT data. Moreover, the EHT model is likely less well specified due to less controlled experimental conditions and may affect the inference of the SB model parameters. The undesirable feedback of information in the inference could be prevented by using the *cut* approach [56] instead of a joint modelling approach. However, we still opt for the joint modelling approach due to the computational burden and the conceptual difficulties of a fully Bayesian interpretation of the cut posterior [57, 58]. We confirmed that the joint modelling approach is sensible by comparing estimates of the resistance characterisation parameters under the joint model with estimates under an SB-only model, as well as by cross-validation in which we predict EHT mortality from ID-SB data only.

In further research, we plan to use the proposed resistance characterisation to better quantify the effectiveness of ITNs over their useful life by taking into account the decay of insecticide concentrations [59]. A possible correlation between insecticide susceptibility and insecticide exposure of individual mosquitoes could be readily accounted for in our model, but the present data do not allow us to identify the needed correlation coefficient. Video tracking data of mosquito-net interaction in cone assays [44] and EHT [45] could be used to estimate this correlation and thus to investigate and isolate the impact of behavioural resistance.

Using a data set from Burkina Faso comprising concurrent ID-SB and EHT studies, our mechanistic model was able to estimate a novel IR metric, including a measure of heterogeneity, from ID-SB data and use this metric to predict ITN killing effects from ID-SB data alone, without requiring EHT data. More paired EHT and ID-SB data from a diverse range of settings, ideally with ID-SB dose escalation reaching approximately LD80, are needed to assess how well our approach generalises across settings. This would enable a larger-scale calibration of transmission models to local insecticide resistance profiles. This is an important step towards refining the recommendations for ITN deployment strategies in the presence of resistance to pyrethroids [9] in order to improve the epidemiological benefit of these important global health tools.

## Acknowledgements

We thank all the researchers who made their data available for inclusion in this study and the anonymous reviewers who have helped to improve the manuscript substantially.

## Supporting information

**S1 Fig. Illustrative example of identifying the parameters that characterise a given resistance setting and predicting the corresponding ITN killing effect with the proposed mechanistic model A** Example killing probabilities for different doses (vertical lines) tested in an intensity-dose susceptibility bioassay (ID-SB) with assumed example dose-response relation (black curve). ‘Dose’ refers to multiples of the discriminating dose defined by the WHO [35]. **B** Contour lines for example killing probabilities (see panel A) in the parameter space characterising the resistance setting. The two-parameter resistance characterisation (black circle), which completely defines the dose-response curve (see panel A), is identifiable if the ID-SB explored the full mortality range across the given doses. **C** Estimated resistance characterisation (white dot) and binned ITN killing probabilities across the parameters space characterising possible resistance characterisations for example parameters of insecticide exposure in EHTs. The parameters of insecticide exposure in EHTs (*µ*_EHT_ and *σ*_EHT_, see Table 2) are identifiable if the data set comprises a sufficient range of ITN killing probabilities. **D** Estimated ITN killing probability for the given example. For this example resistance setting, we arbitrarily choose a median lethal dose of 2.23 (*µ*_T_ = 0.8) and a tolerance heterogeneity measure (standard deviation of the logarithm of the lethal dose, *σ*_T_) of 1. For exposure in SB and EHT assays, we arbitrarily choose the parameter values *σ*_SB_ = 0.2, *µ*_EHT_ = 1.0 and *σ*_EHT_ = 0.5 (see Table 2 for a description of these parameters). For the estimates in the main text, all these parameters were fitted to data.

**S2 Fig. Prior predictive checks.** Mortality in **A** intensity-dose susceptibility bioassays (ID-SB; on the logarithmic scale), **B** discriminating-dose susceptibility bioassays (DD-SB) and **C** experimental hut trials (EHT) under the respective mechanistic models with parameters sampled from the priors (see Table 2). For the ID-SB model, the specified priors support a wide variety of dose-response curves (fine blue curves). The prior predictive mortalities for control SB and DD-SB are identical to those of the corresponding doses in ID-SB.

**S3 Fig. Estimated insecticide resistance characterisation and ITN killing effect. A** Characterisations of insecticide resistance by median lethal dose and proposed measure of heterogeneity in insecticide tolerance (standard deviation of the logarithm of the lethal dose, *σ*_T_) for all assay pairs (see Table 1 for assay pair specification), displayed by the posterior median (white dots). Note that the uncertainty may be very large and the posterior correlation between *µ*_T_ and *σ*_T_ very strong for assay pairs with only DD-SB data (see S9 Table). The coloured areas display binned posterior predictive ITN killing probabilities (posterior median; see panel B for the colour legend) for a range of insecticide resistance characterisations, as estimated by the joint mechanistic model. The red vertical line marks the estimated median insecticide exposure in an EHT (posterior median). **B** Posterior predictive ITN killing probabilities for all assay pairs (see Table 1 for assay pair specification), displayed as posterior median (square white markers) and quantile-based 95% credible intervals (vertical white lines).

**S4 Fig. Leave-one-assay-pair-out cross-validation for assay pairs comprising intensity-dose susceptibility bioassays.** Mosquito mortality in EHT is predicted from ID-SB data only, for each assay pair comprising ID-SBs, using the joint model calibrated to all SB and EHT data except the EHT intervention arm of the given assay pair. These predictions correspond to the estimates in Fig 4 **B**. See Fig 4 for a description of the plot elements.

**S5 Fig. Simulation study on the sufficiency of dose escalation for estimation of the lethal dose distribution. A** Mosquito mortality probability in ID-SBs under increasing insecticide challenge (black curve) and simulated mosquito death proportions (black dots, plotted with random jitter) for nine potential resistance metrics, under the SB model with calibrated exposure parameters. The three *µ_T_* values (mean of log lethal dose) approximately correspond to the lowest and highest values estimated across assay pairs 2, 5, 6 and 7, plus double the highest value. The three *σ_T_*values (standard deviation of log lethal dose) approximately correspond to the lowest and highest values estimated across assay pairs 1–7, plus a middle value. We set the standard deviation of the log insecticide exposure and the control mortality probability to the corresponding posterior medians from the SB fit (0.1293 and 0.016, respectively, see S8 Table). For each dose, we simulated five replicates of death counts using a beta-binomial model (mean–dispersion parameterisation) with a total count of 50 for each and an intraclass correlation coefficient of 0.1, and then computed proportions. **B** Given lethal dose distribution (thick black) and corresponding estimates based on the simulated ID-SB data (panel A) including the control, the discriminating dose and dose-escalation from LD10 up to the indicated LD level (coloured curves).

**S6 Table. Summary of the paired DD-SB and EHT data used in this study.** Note that controls may be shared between assay pairs. For a summary of the paired ID-SB and EHT data used in this study see Table 1.

References correspond to the bibliography in the main text.

**S7 Table. Parameter estimates and sampling diagnostics from the joint SB and EHT model** Mean, median, standard deviation (SD), median absolute deviation (MAD), 5% Quantile, 95% Quantile, a MCMC convergence diagnostic (Rhat), bulk effective sample size (ESS bulk) and tail effective sample size (ESS tail) for all posterior parameter samples of the joint model for SB and EHT data. For explanations on MCMC inference diagnostics, see [53].

**S8 Table. Parameter estimates and sampling diagnostics from the SB only model** Mean, median, standard deviation (SD), median absolute deviation (MAD), 5% Quantile, 95% Quantile, a MCMC convergence diagnostic (Rhat), bulk effective sample size (ESS bulk) and tail effective sample size (ESS tail) for all posterior parameter samples of the model for SB data only. For explanations on MCMC inference diagnostics, see [53].

**S9 Table. Model predictions per assay pair from the joint SB and EHT model** Point estimates (median) and interval estimates (quantile-based 95% credible intervals) for the estimated ITN killing effect, the median lethal dose (LD_50_) and the standard deviation of the logarithm of the lethal dose (*σ*_T_) as a measure of heterogeneity in insecticide tolerance, as estimated from the joint SB and EHT model. The Pearson correlation coefficient (Cor(*µ*_T_*, σ*_T_)) between the mean (*µ*_T_) and the standard deviation (*σ*_T_) of the logarithm of the lethal dose indicates how well the resistance characterisation by these two parameters could be identified from the data. The closer 0 is, the better the characterisation could be identified.

