## Supplementary figures and images for "Inferring a novel insecticide resistance metric and exposure variability in mosquito bioassays across Africa"

### S1 Fig

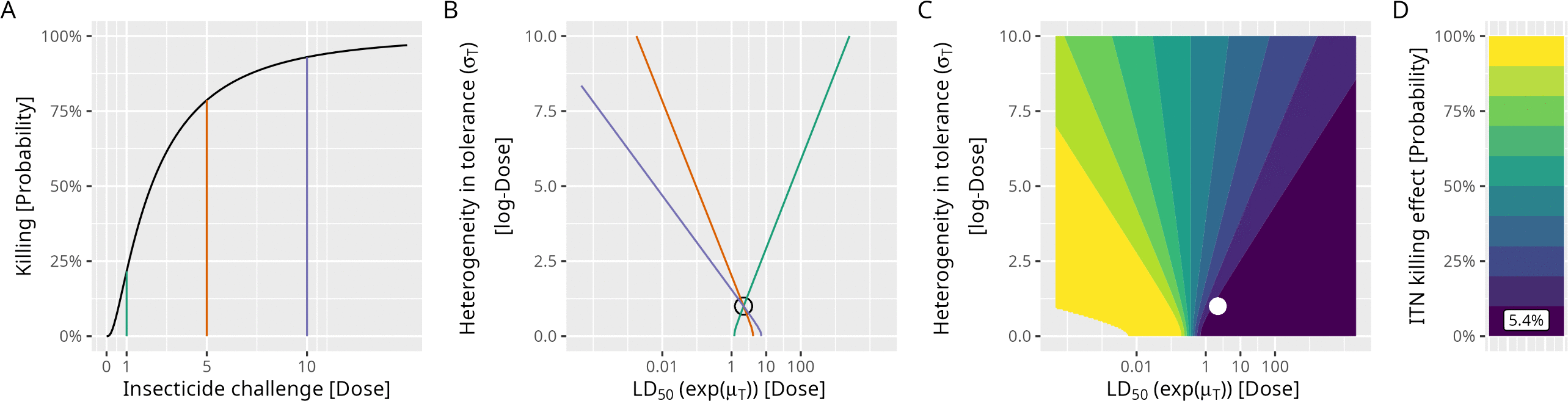

### S2 Fig

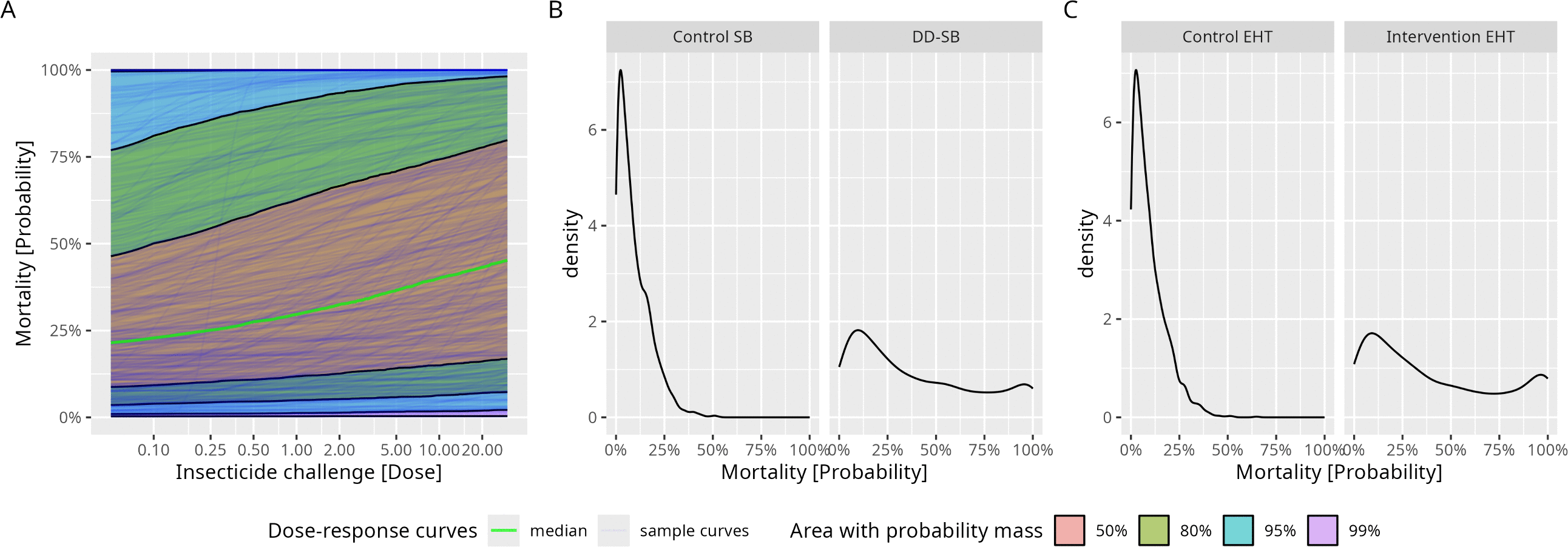

### S3 Fig

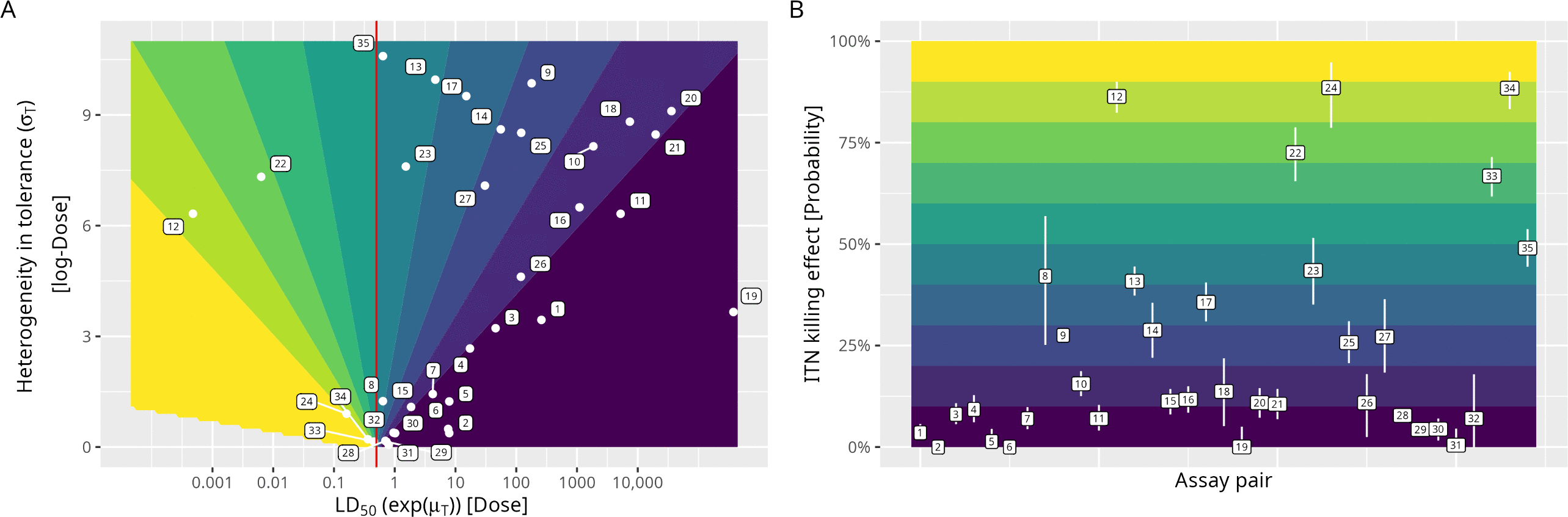

### S4 Fig

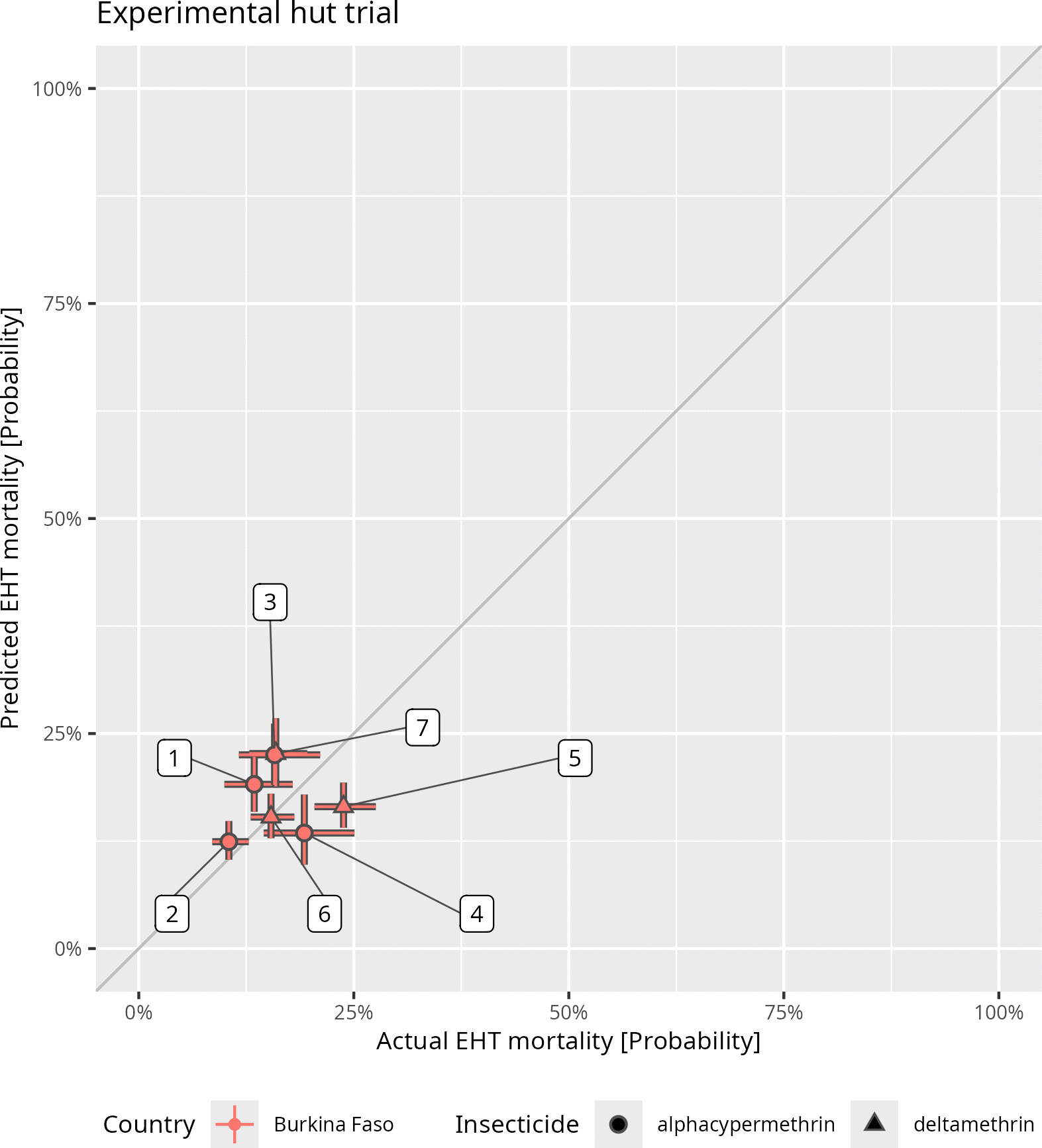

### S5 Fig

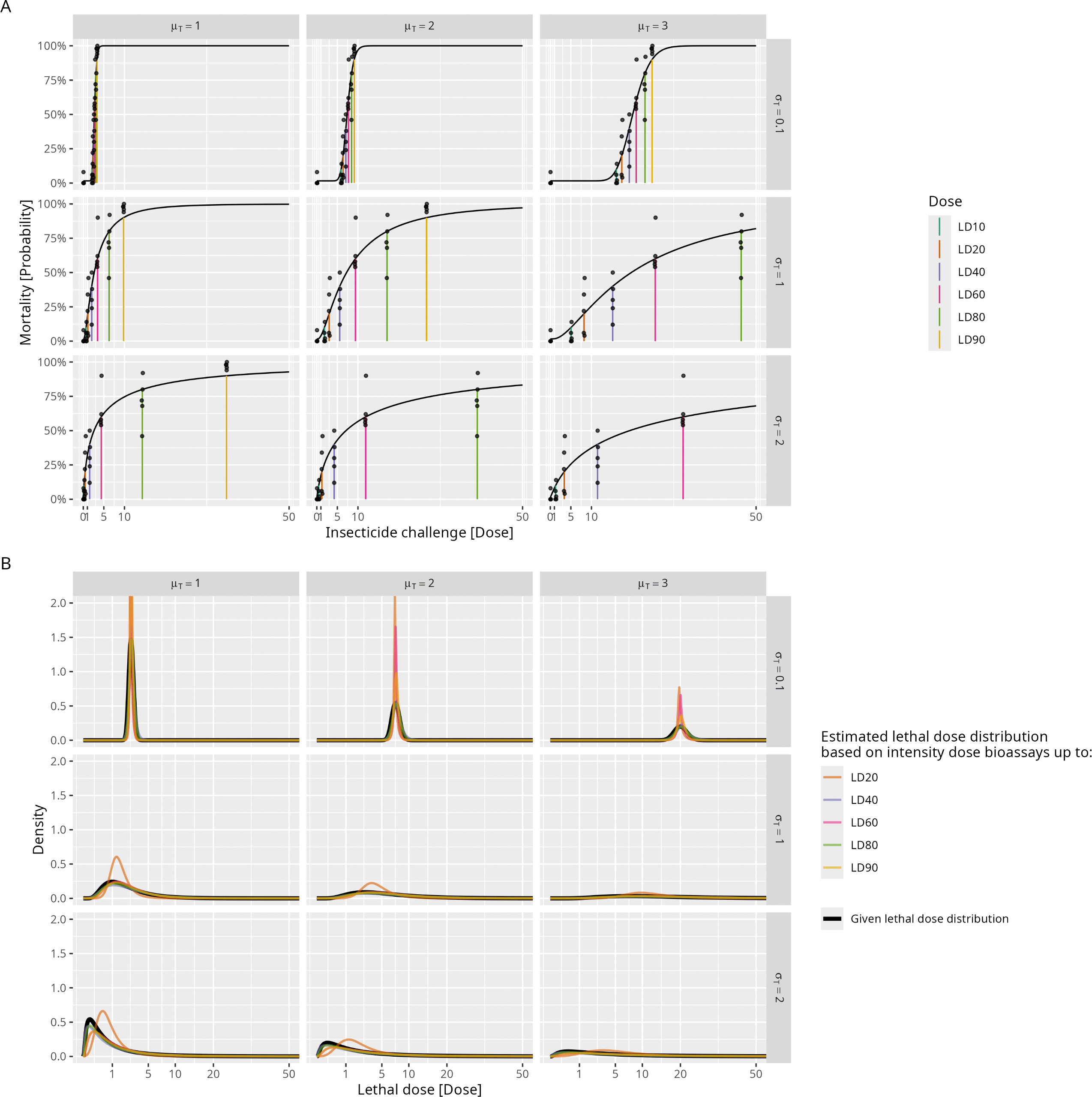
