## Supplementary material for "Inferring a novel insecticide resistance metric and exposure variability in mosquito bioassays across Africa": S6 Table

### Summary of the paired DD-SB and EHT data used in this study.

Note that controls may be shared between assay pairs. For a summary of the paired ID-SB and EHT data used in this study see Table 1. References correspond to the bibliography in the main text.

| Assay pair | Reference | Country | Site | Year | SB protocol | Hut type | Insecticides in SB | Insecticides on ITNs | ITN products | Total in treatment SB | Total in control SB | Total in intervention hut | Total in control hut |
|---|---|---|---|---|---|---|---|---|---|---|---|---|---|
| 8 | [60] | Benin | Akron | 2010 | WHO | West | deltamethrin | deltamethrin | LifeNet | 61 | 0 | 63 | 144 |
| 9 | [61] | Benin | Cove | 2010 | WHO | West | permethrin | permethrin | Olyset | 100 | 0 | 3804 | 2874 |
| 10 | [62] | Benin | Cove | 2012 | WHO | West | deltamethrin, permethrin | alphacypermethrin | Interceptor | 100 | 0 | 631 | 673 |
| 11 | [63] | Benin | Cove | 2015 | WHO | West | deltamethrin, permethrin | alphacypermethrin | Interceptor | 200 | 0 | 175 | 310 |
| 12 | [64] | Benin | Malanville | 2008 | WHO | West | deltamethrin | deltamethrin | PermaNet2 | 100 | 0 | 243 | 285 |
| 13 | [60] | Benin | Malanville | 2010 | WHO | West | deltamethrin | deltamethrin | LifeNet | 97 | 0 | 676 | 911 |
| 14 | [65] | Benin | Malanville | 2011 | WHO | West | permethrin | permethrin | Olyset | 100 | 0 | 96 | 69 |
| 15 | [66] | Burkina Faso | Tengrela | 2014 | WHO | West | deltamethrin | deltamethrin | DawaPlus2, PermaNet2 | 85 | 0 | 1485 | 482 |
| 16 | [66] | Burkina Faso | Tengrela | 2014 | WHO | West | permethrin | permethrin | Olyset | 101 | 0 | 922 | 482 |
| 17 | [64] | Burkina Faso | VDK | 2007 | WHO | West | deltamethrin | deltamethrin | PermaNet2 | 100 | 0 | 329 | 908 |
| 18 | [67] | Burkina Faso | VDK | 2011 | WHO | West | deltamethrin | deltamethrin | PermaNet2 | 100 | 0 | 114 | 336 |
| 19 | [68] | Burkina Faso | VDK | 2014 | WHO | West | deltamethrin, permethrin | alphacypermethrin | Interceptor | 227 | 0 | 519 | 853 |
| 20 | [66] | Burkina Faso | VK5 | 2014 | WHO | West | deltamethrin | deltamethrin | DawaPlus2, PermaNet2 | 163 | 0 | 2532 | 1095 |
| 21 | [66] | Burkina Faso | VK5 | 2014 | WHO | West | permethrin | permethrin | Olyset | 153 | 0 | 1458 | 1095 |
| 22 | [64] | Cameroon | Pitoa | 2008 | WHO | West | deltamethrin | deltamethrin | PermaNet2 | 100 | 0 | 105 | 401 |
| 23 | [69] | Cote d'Ivoire | Tiassale | 2012 | WHO | West | deltamethrin | deltamethrin | PermaNet2 | 99 | 0 | 108 | 130 |
| 24 | [70] | Cote d'Ivoire | Yaokoffikro | 2000 | pre2000 | West | pyrethroid | deltamethrin | PermaNet | 50 | 0 | 69 | 84 |
| 25 | [71] | Cote d'Ivoire | Yaokoffikro | 2009 | WHO | West | deltamethrin | deltamethrin | PermaNet2 | 50 | 0 | 317 | 796 |
| 26 | [72] | Ethiopia | GGDam | 2011 | WHO | East | deltamethrin | deltamethrin | PermaNet2 | 80 | 0 | 426 | 511 |
| 27 | [73] | Tanzania | LowerMoshi | 2013 | WHO | East | deltamethrin | deltamethrin | PermaNet2 | 100 | 0 | 10 | 75 |
| 28 | [74] | Tanzania | Lupiro | 2010 | WHO | Ifakara | deltamethrin | deltamethrin | PermaNet2, Icon Life | 96 | 0 | 30431 | 11777 |
| 29 | [74] | Tanzania | Lupiro | 2010 | WHO | Ifakara | permethrin | permethrin | Olyset | 125 | 0 | 15836 | 11777 |
| 30 | [75] | Tanzania | Lupiro | 2015 | WHO | Ifakara | deltamethrin, lambdacyhalothrin, permethrin | alphacypermethrin | MAGNet | 150 | 0 | 1070 | 810 |
| 31 | [76] | Tanzania | Mabogini | 2005 | WHO | East | permethrin | permethrin | Olyset | 100 | 0 | 196 | 716 |
| 32 | [75] | Tanzania | Muheza | 2015 | WHO | East | permethrin | alphacypermethrin | Interceptor | 50 | 0 | 94 | 139 |
| 33 | [77] | Tanzania | Zeneti | 2005 | WHO | East | permethrin | permethrin | Olyset | 393 | 0 | 399 | 854 |
| 34 | [78] | Tanzania | Zeneti | 2006 | WHO | East | alphacypermethrin | alphacypermethrin | Interceptor | 752 | 0 | 202 | 252 |
| 35 | [79] | Togo | Kolokope | 2013 | WHO | West | deltamethrin | deltamethrin | Yorkool | 74 | 0 | 389 | 465 |
