## Supplementary material for "Inferring a novel insecticide resistance metric and exposure variability in mosquito bioassays across Africa": S7 Table: S7_Table.html

|  |  |  |  |  |  |  |  |  |  |  |
| --- | --- | --- | --- | --- | --- | --- | --- | --- | --- | --- |
| Parameter estimates and sampling diagnostics | | | | | | | | | | |
| from joint SB and EHT model | | | | | | | | | | |
| Parameter | Assay pair | Mean | Median | SD^1^ | MAD^2^ | 5% Quantile | 95% Quantile | Rhat^3^ | ESS\_bulk^4^ | ESS\_tail^5^ |
| p^(0)^_SB_ | - | 0.01454 | 0.01451 | 0.001300 | 0.001276 | 0.01246 | 0.01673 | 1.000 | 14,070 | 8,347 |
| σ_V_ | - | 0.1124 | 0.1067 | 0.06906 | 0.07167 | 0.01378 | 0.2374 | 1.001 | 1,660 | 1,242 |
| μ_X_ | - | −0.7014 | −0.6929 | 0.1405 | 0.1382 | −0.9513 | −0.4860 | 1.001 | 1,870 | 1,560 |
| σ_X_ | - | 0.1482 | 0.1433 | 0.08146 | 0.08216 | 0.02121 | 0.2896 | 1.001 | 3,066 | 3,468 |
| μ_T_ | 1 | 5.671 | 5.559 | 0.8091 | 0.7358 | 4.592 | 7.170 | 1.000 | 6,628 | 5,162 |
| μ_T_ | 2 | 2.067 | 2.066 | 0.03435 | 0.03407 | 2.011 | 2.124 | 1.000 | 16,490 | 8,577 |
| μ_T_ | 3 | 3.854 | 3.823 | 0.3142 | 0.2990 | 3.394 | 4.414 | 1.001 | 8,195 | 5,620 |
| μ_T_ | 4 | 2.870 | 2.851 | 0.1895 | 0.1773 | 2.587 | 3.212 | 1.000 | 8,865 | 6,425 |
| μ_T_ | 5 | 2.081 | 2.068 | 0.1394 | 0.1326 | 1.875 | 2.332 | 1.000 | 9,989 | 6,506 |
| μ_T_ | 6 | 2.024 | 2.024 | 0.02451 | 0.02429 | 1.983 | 2.064 | 1.000 | 14,540 | 7,303 |
| μ_T_ | 7 | 1.443 | 1.443 | 0.04994 | 0.05043 | 1.363 | 1.526 | 1.000 | 15,240 | 9,413 |
| μ_T_ | 8 | −0.5302 | −0.4477 | 0.4027 | 0.2149 | −1.213 | −0.1595 | 1.001 | 4,502 | 2,885 |
| μ_T_ | 9 | 5.354 | 5.188 | 1.741 | 1.758 | 2.791 | 8.429 | 1.000 | 7,138 | 6,153 |
| μ_T_ | 10 | 7.828 | 7.528 | 2.883 | 2.916 | 3.651 | 12.96 | 1.000 | 7,768 | 7,200 |
| μ_T_ | 11 | 9.036 | 8.562 | 3.807 | 3.797 | 3.666 | 15.96 | 1.000 | 6,756 | 7,793 |
| μ_T_ | 12 | −7.983 | −7.647 | 2.844 | 2.807 | −13.24 | −3.946 | 1.001 | 7,427 | 7,665 |
| μ_T_ | 13 | 1.650 | 1.536 | 0.8373 | 0.8045 | 0.4725 | 3.173 | 1.000 | 8,961 | 6,950 |
| μ_T_ | 14 | 4.255 | 4.021 | 1.757 | 1.672 | 1.826 | 7.514 | 1.000 | 8,121 | 7,030 |
| μ_T_ | 15 | 0.8415 | 0.6208 | 0.8417 | 0.3708 | 0.2270 | 2.118 | 1.001 | 3,648 | 1,898 |
| μ_T_ | 16 | 7.378 | 7.003 | 3.230 | 3.217 | 2.859 | 13.20 | 1.000 | 7,288 | 7,902 |
| μ_T_ | 17 | 2.880 | 2.712 | 1.250 | 1.188 | 1.174 | 5.181 | 1.000 | 8,498 | 6,209 |
| μ_T_ | 18 | 9.283 | 8.911 | 3.257 | 3.186 | 4.683 | 15.15 | 1.000 | 8,142 | 7,385 |
| μ_T_ | 19 | 13.50 | 12.84 | 6.032 | 5.750 | 4.764 | 24.48 | 1.000 | 8,716 | 6,738 |
| μ_T_ | 20 | 10.80 | 10.49 | 3.331 | 3.315 | 5.897 | 16.77 | 1.000 | 6,243 | 7,287 |
| μ_T_ | 21 | 10.16 | 9.889 | 3.259 | 3.291 | 5.319 | 15.88 | 1.001 | 7,007 | 8,143 |
| μ_T_ | 22 | −5.330 | −5.061 | 2.003 | 1.918 | −9.036 | −2.568 | 1.000 | 7,119 | 7,027 |
| μ_T_ | 23 | 0.5354 | 0.4212 | 0.9242 | 0.7822 | −0.7704 | 2.210 | 1.000 | 8,594 | 6,689 |
| μ_T_ | 24 | −2.928 | −1.827 | 2.673 | 1.127 | −8.797 | −0.8601 | 1.003 | 3,380 | 4,348 |
| μ_T_ | 25 | 5.022 | 4.794 | 2.009 | 1.960 | 2.152 | 8.703 | 1.000 | 7,903 | 7,468 |
| μ_T_ | 26 | 5.343 | 4.782 | 3.366 | 3.400 | 0.9395 | 11.64 | 1.001 | 5,955 | 6,263 |
| μ_T_ | 27 | 3.713 | 3.418 | 1.841 | 1.735 | 1.253 | 7.159 | 1.000 | 8,623 | 7,598 |
| μ_T_ | 28 | −0.3636 | −0.3554 | 0.09506 | 0.09103 | −0.5332 | −0.2258 | 1.002 | 1,607 | 1,261 |
| μ_T_ | 29 | −0.3204 | −0.3122 | 0.09460 | 0.09094 | −0.4859 | −0.1794 | 1.002 | 1,650 | 1,166 |
| μ_T_ | 30 | 0.01795 | 0.01426 | 0.04496 | 0.04103 | −0.04846 | 0.09763 | 1.001 | 11,410 | 5,273 |
| μ_T_ | 31 | −0.2380 | −0.2278 | 0.1029 | 0.1001 | −0.4212 | −0.08842 | 1.001 | 2,028 | 1,226 |
| μ_T_ | 32 | −0.04089 | −0.04451 | 0.08546 | 0.06770 | −0.1670 | 0.1024 | 1.003 | 7,356 | 4,918 |
| μ_T_ | 33 | −0.8008 | −0.7904 | 0.1642 | 0.1625 | −1.089 | −0.5526 | 1.001 | 2,063 | 2,748 |
| μ_T_ | 34 | −1.045 | −1.019 | 0.2538 | 0.2419 | −1.499 | −0.6794 | 1.001 | 2,719 | 4,873 |
| μ_T_ | 35 | −0.4364 | −0.4457 | 0.6928 | 0.6303 | −1.539 | 0.6957 | 1.000 | 10,700 | 7,726 |
| σ_T_ | 1 | 3.541 | 3.446 | 0.6683 | 0.6122 | 2.640 | 4.761 | 1.000 | 6,730 | 5,295 |
| σ_T_ | 2 | 0.3749 | 0.3778 | 0.05109 | 0.04013 | 0.2994 | 0.4461 | 1.001 | 2,470 | 1,280 |
| σ_T_ | 3 | 3.259 | 3.221 | 0.3802 | 0.3631 | 2.707 | 3.947 | 1.000 | 8,001 | 5,740 |
| σ_T_ | 4 | 2.697 | 2.671 | 0.2912 | 0.2774 | 2.265 | 3.216 | 1.000 | 7,912 | 6,851 |
| σ_T_ | 5 | 1.252 | 1.236 | 0.2000 | 0.1946 | 0.9499 | 1.602 | 1.000 | 7,561 | 7,559 |
| σ_T_ | 6 | 0.4912 | 0.4906 | 0.04621 | 0.04107 | 0.4200 | 0.5650 | 1.001 | 3,617 | 1,691 |
| σ_T_ | 7 | 1.440 | 1.437 | 0.07163 | 0.07142 | 1.327 | 1.563 | 1.000 | 13,350 | 9,139 |
| σ_T_ | 8 | 1.993 | 1.245 | 1.997 | 0.7479 | 0.5511 | 6.354 | 1.001 | 3,322 | 4,498 |
| σ_T_ | 9 | 10.15 | 9.857 | 2.936 | 2.985 | 5.794 | 15.34 | 1.001 | 6,503 | 6,660 |
| σ_T_ | 10 | 8.425 | 8.149 | 2.844 | 2.865 | 4.271 | 13.52 | 1.000 | 7,650 | 7,606 |
| σ_T_ | 11 | 6.637 | 6.321 | 2.686 | 2.721 | 2.798 | 11.54 | 1.000 | 6,699 | 7,893 |
| σ_T_ | 12 | 6.603 | 6.324 | 2.461 | 2.469 | 3.104 | 11.16 | 1.001 | 7,546 | 7,655 |
| σ_T_ | 13 | 10.19 | 9.956 | 3.055 | 3.111 | 5.617 | 15.64 | 1.000 | 7,482 | 7,059 |
| σ_T_ | 14 | 8.900 | 8.611 | 2.951 | 2.932 | 4.572 | 14.25 | 1.000 | 7,550 | 8,112 |
| σ_T_ | 15 | 1.276 | 1.087 | 0.7909 | 0.3939 | 0.6259 | 2.484 | 1.001 | 2,653 | 1,979 |
| σ_T_ | 16 | 6.809 | 6.497 | 2.758 | 2.763 | 2.864 | 11.78 | 1.000 | 7,260 | 7,994 |
| σ_T_ | 17 | 9.788 | 9.511 | 3.046 | 2.990 | 5.297 | 15.19 | 1.001 | 7,234 | 7,163 |
| σ_T_ | 18 | 9.031 | 8.817 | 2.854 | 2.812 | 4.768 | 14.09 | 1.000 | 8,058 | 8,034 |
| σ_T_ | 19 | 4.157 | 3.660 | 3.018 | 3.374 | 0.3295 | 9.640 | 1.001 | 4,415 | 6,707 |
| σ_T_ | 20 | 9.328 | 9.105 | 2.729 | 2.732 | 5.285 | 14.16 | 1.000 | 6,209 | 7,505 |
| σ_T_ | 21 | 8.682 | 8.471 | 2.660 | 2.732 | 4.696 | 13.30 | 1.001 | 7,149 | 8,155 |
| σ_T_ | 22 | 7.649 | 7.325 | 2.810 | 2.780 | 3.624 | 12.74 | 1.000 | 6,946 | 7,523 |
| σ_T_ | 23 | 7.924 | 7.602 | 3.062 | 3.109 | 3.520 | 13.45 | 1.000 | 8,609 | 7,162 |
| σ_T_ | 24 | 1.703 | 0.9023 | 1.956 | 0.8618 | 0.1498 | 5.990 | 1.002 | 3,627 | 4,510 |
| σ_T_ | 25 | 8.791 | 8.516 | 2.957 | 2.972 | 4.375 | 14.11 | 1.000 | 7,425 | 7,635 |
| σ_T_ | 26 | 5.050 | 4.615 | 3.008 | 3.101 | 0.9886 | 10.63 | 1.001 | 5,899 | 5,899 |
| σ_T_ | 27 | 7.441 | 7.086 | 3.078 | 3.108 | 3.044 | 13.04 | 1.000 | 7,698 | 7,769 |
| σ_T_ | 28 | 0.1698 | 0.1676 | 0.05421 | 0.05136 | 0.08533 | 0.2615 | 1.002 | 2,187 | 2,230 |
| σ_T_ | 29 | 0.1445 | 0.1457 | 0.05580 | 0.05275 | 0.04610 | 0.2346 | 1.002 | 1,930 | 1,485 |
| σ_T_ | 30 | 0.3837 | 0.3764 | 0.1008 | 0.09568 | 0.2329 | 0.5612 | 1.001 | 2,102 | 2,560 |
| σ_T_ | 31 | 0.06443 | 0.05671 | 0.04612 | 0.04787 | 0.005294 | 0.1515 | 1.000 | 4,633 | 4,578 |
| σ_T_ | 32 | 0.4054 | 0.3890 | 0.1909 | 0.1653 | 0.1084 | 0.7330 | 1.002 | 2,607 | 2,139 |
| σ_T_ | 33 | 0.1527 | 0.1476 | 0.09056 | 0.09749 | 0.01807 | 0.3111 | 0.9999 | 4,354 | 3,991 |
| σ_T_ | 34 | 0.2242 | 0.2129 | 0.1331 | 0.1384 | 0.02946 | 0.4562 | 1.001 | 5,013 | 3,907 |
| σ_T_ | 35 | 10.86 | 10.59 | 3.061 | 3.042 | 6.281 | 16.31 | 1.001 | 11,110 | 8,620 |
| p^(0)^_EHT_ | 1 | 0.1417 | 0.1415 | 0.01121 | 0.01110 | 0.1235 | 0.1607 | 1.001 | 18,240 | 9,462 |
| p^(0)^_EHT_ | 2 | 0.1154 | 0.1152 | 0.007720 | 0.007777 | 0.1030 | 0.1282 | 1.000 | 16,130 | 9,614 |
| p^(0)^_EHT_ | 3 | 0.1360 | 0.1358 | 0.01224 | 0.01233 | 0.1162 | 0.1567 | 1.001 | 14,490 | 8,327 |
| p^(0)^_EHT_ | 4 | 0.07560 | 0.07478 | 0.01433 | 0.01445 | 0.05345 | 0.1002 | 1.001 | 13,650 | 8,782 |
| p^(0)^_EHT_ | 5 | 0.1843 | 0.1841 | 0.01150 | 0.01147 | 0.1654 | 0.2034 | 0.9999 | 13,000 | 9,172 |
| p^(0)^_EHT_ | 6 | 0.1537 | 0.1536 | 0.009159 | 0.009220 | 0.1388 | 0.1688 | 1.000 | 16,150 | 7,723 |
| p^(0)^_EHT_ | 7 | 0.1315 | 0.1313 | 0.01146 | 0.01149 | 0.1128 | 0.1507 | 0.9999 | 11,090 | 8,992 |
| p^(0)^_EHT_ | 8 | 0.1487 | 0.1470 | 0.02811 | 0.02823 | 0.1056 | 0.1973 | 1.001 | 14,770 | 8,682 |
| p^(0)^_EHT_ | 9 | 0.05693 | 0.05685 | 0.004398 | 0.004458 | 0.04992 | 0.06430 | 0.9999 | 16,710 | 9,152 |
| p^(0)^_EHT_ | 10 | 0.03196 | 0.03154 | 0.006933 | 0.006699 | 0.02144 | 0.04403 | 1.000 | 14,840 | 7,904 |
| p^(0)^_EHT_ | 11 | 0.03068 | 0.02972 | 0.009764 | 0.009689 | 0.01661 | 0.04808 | 1.000 | 14,480 | 7,783 |
| p^(0)^_EHT_ | 12 | 0.04499 | 0.04404 | 0.01217 | 0.01185 | 0.02669 | 0.06653 | 1.000 | 15,410 | 7,694 |
| p^(0)^_EHT_ | 13 | 0.01831 | 0.01792 | 0.004605 | 0.004551 | 0.01150 | 0.02658 | 0.9999 | 16,940 | 8,159 |
| p^(0)^_EHT_ | 14 | 0.02236 | 0.01644 | 0.02082 | 0.01644 | 0.001300 | 0.06442 | 1.001 | 9,710 | 5,008 |
| p^(0)^_EHT_ | 15 | 0.07842 | 0.07768 | 0.01204 | 0.01191 | 0.05957 | 0.09948 | 1.000 | 11,870 | 8,824 |
| p^(0)^_EHT_ | 16 | 0.08381 | 0.08341 | 0.01225 | 0.01228 | 0.06452 | 0.1048 | 1.001 | 14,970 | 7,644 |
| p^(0)^_EHT_ | 17 | 0.05163 | 0.05136 | 0.007458 | 0.007336 | 0.03985 | 0.06449 | 1.000 | 16,170 | 8,898 |
| p^(0)^_EHT_ | 18 | 0.1510 | 0.1502 | 0.02317 | 0.02309 | 0.1142 | 0.1905 | 1.000 | 13,400 | 9,320 |
| p^(0)^_EHT_ | 19 | 0.1364 | 0.1368 | 0.01147 | 0.01148 | 0.1169 | 0.1544 | 1.000 | 8,064 | 7,593 |
| p^(0)^_EHT_ | 20 | 0.2154 | 0.2153 | 0.01322 | 0.01325 | 0.1937 | 0.2371 | 1.001 | 15,680 | 8,943 |
| p^(0)^_EHT_ | 21 | 0.2060 | 0.2058 | 0.01239 | 0.01237 | 0.1859 | 0.2265 | 1.001 | 14,000 | 8,718 |
| p^(0)^_EHT_ | 22 | 0.1315 | 0.1309 | 0.01677 | 0.01661 | 0.1050 | 0.1604 | 0.9999 | 15,690 | 9,392 |
| p^(0)^_EHT_ | 23 | 0.2087 | 0.2075 | 0.03421 | 0.03414 | 0.1550 | 0.2667 | 1.000 | 15,100 | 8,867 |
| p^(0)^_EHT_ | 24 | 0.09273 | 0.08977 | 0.02939 | 0.02907 | 0.05018 | 0.1454 | 1.000 | 14,540 | 7,068 |
| p^(0)^_EHT_ | 25 | 0.07171 | 0.07129 | 0.009303 | 0.009010 | 0.05696 | 0.08778 | 1.000 | 16,610 | 8,400 |
| p^(0)^_EHT_ | 26 | 0.5506 | 0.5508 | 0.01823 | 0.01803 | 0.5208 | 0.5803 | 1.000 | 13,040 | 7,644 |
| p^(0)^_EHT_ | 27 | 0.02658 | 0.02250 | 0.01819 | 0.01598 | 0.004697 | 0.06155 | 1.000 | 13,090 | 6,053 |
| p^(0)^_EHT_ | 28 | 0.1165 | 0.1164 | 0.002947 | 0.002942 | 0.1117 | 0.1214 | 1.000 | 17,120 | 9,106 |
| p^(0)^_EHT_ | 29 | 0.1165 | 0.1166 | 0.002932 | 0.002976 | 0.1117 | 0.1214 | 1.000 | 17,970 | 8,924 |
| p^(0)^_EHT_ | 30 | 0.05991 | 0.05946 | 0.008580 | 0.008388 | 0.04652 | 0.07492 | 1.000 | 13,030 | 7,464 |
| p^(0)^_EHT_ | 31 | 0.1689 | 0.1688 | 0.01237 | 0.01223 | 0.1489 | 0.1901 | 1.001 | 15,730 | 8,914 |
| p^(0)^_EHT_ | 32 | 0.09275 | 0.09128 | 0.02289 | 0.02304 | 0.05790 | 0.1329 | 1.000 | 9,159 | 8,507 |
| p^(0)^_EHT_ | 33 | 0.06833 | 0.06797 | 0.008542 | 0.008527 | 0.05480 | 0.08284 | 1.000 | 18,590 | 8,925 |
| p^(0)^_EHT_ | 34 | 0.09127 | 0.09009 | 0.01769 | 0.01742 | 0.06422 | 0.1221 | 1.001 | 15,450 | 9,084 |
| p^(0)^_EHT_ | 35 | 0.02393 | 0.02316 | 0.007518 | 0.007305 | 0.01315 | 0.03721 | 1.000 | 16,240 | 8,626 |
|  |  |  |  |  |  |  |  |  |  |  |
| --- | --- | --- | --- | --- | --- | --- | --- | --- | --- | --- |
| Reference for sampling diagnostics: Aki Vehtari, Andrew Gelman, Daniel Simpson, Bob Carpenter, and Paul-Christian Bürkner (2021). Rank-normalization, folding, and localization: An improved R-hat for assessing convergence of MCMC (with discussion). *Bayesian Data Analysis*. 16(2), 667-–718. doi:10.1214/20-BA1221 | | | | | | | | | | |
|  |  |  |  |  |  |  |  |  |  |  |
| --- | --- | --- | --- | --- | --- | --- | --- | --- | --- | --- |
| ^1^ Standard deviation | | | | | | | | | | |
| ^2^ Median absolute deviation | | | | | | | | | | |
| ^3^ Rhat MCMC convergence diagnostic | | | | | | | | | | |
| ^4^ Bulk effective sample size | | | | | | | | | | |
| ^5^ Tail effective sample size | | | | | | | | | | |
