## Supplementary material for "Inferring a novel insecticide resistance metric and exposure variability in mosquito bioassays across Africa": S8 Table: S8_Table.html

|  |  |  |  |  |  |  |  |  |  |  |
| --- | --- | --- | --- | --- | --- | --- | --- | --- | --- | --- |
| Parameter estimates and sampling diagnostics | | | | | | | | | | |
| from SB only model | | | | | | | | | | |
| Parameter | Assay pair | Mean | Median | SD^1^ | MAD^2^ | 5% Quantile | 95% Quantile | Rhat^3^ | ESS\_bulk^4^ | ESS\_tail^5^ |
| σ_V_ | - | 0.2865 | 0.3237 | 0.1208 | 0.1099 | 0.04709 | 0.4333 | 1.001 | 2,395 | 2,320 |
| p^(0)^_SB_ | 1 | 0.007098 | 0.006768 | 0.002620 | 0.002517 | 0.003405 | 0.01190 | 1.001 | 10,040 | 5,607 |
| p^(0)^_SB_ | 2 | 0.02480 | 0.02449 | 0.004768 | 0.004750 | 0.01747 | 0.03318 | 1.001 | 9,501 | 6,120 |
| p^(0)^_SB_ | 3 | 0.01381 | 0.01359 | 0.003037 | 0.002975 | 0.009184 | 0.01914 | 1.001 | 10,280 | 5,823 |
| p^(0)^_SB_ | 4 | 0.01409 | 0.01377 | 0.003831 | 0.003773 | 0.008386 | 0.02080 | 1.000 | 10,220 | 5,215 |
| p^(0)^_SB_ | 5 | 0.01139 | 0.009858 | 0.007078 | 0.006327 | 0.002850 | 0.02529 | 0.9998 | 6,448 | 4,708 |
| p^(0)^_SB_ | 6 | 0.02520 | 0.02499 | 0.004107 | 0.004084 | 0.01889 | 0.03235 | 1.001 | 9,525 | 5,335 |
| p^(0)^_SB_ | 7 | 0.01368 | 0.01341 | 0.003169 | 0.003062 | 0.008965 | 0.01930 | 1.000 | 9,106 | 4,970 |
| μ_T_ | 1 | 6.379 | 6.181 | 1.135 | 0.9525 | 4.953 | 8.534 | 1.001 | 5,501 | 3,463 |
| μ_T_ | 2 | 2.077 | 2.076 | 0.03555 | 0.03464 | 2.020 | 2.138 | 1.001 | 9,776 | 5,810 |
| μ_T_ | 3 | 4.050 | 3.999 | 0.3881 | 0.3703 | 3.507 | 4.757 | 1.001 | 4,805 | 3,669 |
| μ_T_ | 4 | 2.809 | 2.794 | 0.1759 | 0.1675 | 2.546 | 3.123 | 1.001 | 5,830 | 4,184 |
| μ_T_ | 5 | 2.044 | 2.037 | 0.1190 | 0.1154 | 1.864 | 2.254 | 1.000 | 7,867 | 5,132 |
| μ_T_ | 6 | 2.033 | 2.033 | 0.02369 | 0.02379 | 1.995 | 2.071 | 1.000 | 8,768 | 6,123 |
| μ_T_ | 7 | 1.428 | 1.428 | 0.05288 | 0.05284 | 1.341 | 1.515 | 1.001 | 10,700 | 6,106 |
| σ_T_ | 1 | 4.325 | 4.147 | 0.9896 | 0.8337 | 3.085 | 6.193 | 1.001 | 5,528 | 3,571 |
| σ_T_ | 2 | 0.2199 | 0.2225 | 0.1264 | 0.1679 | 0.02207 | 0.4043 | 1.001 | 2,436 | 2,675 |
| σ_T_ | 3 | 3.577 | 3.512 | 0.4985 | 0.4710 | 2.877 | 4.507 | 1.001 | 4,807 | 3,704 |
| σ_T_ | 4 | 2.475 | 2.447 | 0.2839 | 0.2704 | 2.062 | 2.983 | 1.001 | 5,700 | 4,419 |
| σ_T_ | 5 | 1.091 | 1.097 | 0.1928 | 0.1685 | 0.7729 | 1.383 | 1.000 | 4,865 | 2,538 |
| σ_T_ | 6 | 0.3737 | 0.3826 | 0.1026 | 0.1049 | 0.1898 | 0.5196 | 1.003 | 2,744 | 1,620 |
| σ_T_ | 7 | 1.478 | 1.476 | 0.08779 | 0.08765 | 1.339 | 1.629 | 1.001 | 8,759 | 5,970 |
|  |  |  |  |  |  |  |  |  |  |  |
| --- | --- | --- | --- | --- | --- | --- | --- | --- | --- | --- |
| Reference for sampling diagnostics: Aki Vehtari, Andrew Gelman, Daniel Simpson, Bob Carpenter, and Paul-Christian Bürkner (2021). Rank-normalization, folding, and localization: An improved R-hat for assessing convergence of MCMC (with discussion). *Bayesian Data Analysis*. 16(2), 667-–718. doi:10.1214/20-BA1221 | | | | | | | | | | |
|  |  |  |  |  |  |  |  |  |  |  |
| --- | --- | --- | --- | --- | --- | --- | --- | --- | --- | --- |
| ^1^ Standard deviation | | | | | | | | | | |
| ^2^ Median absolute deviation | | | | | | | | | | |
| ^3^ Rhat MCMC convergence diagnostic | | | | | | | | | | |
| ^4^ Bulk effective sample size | | | | | | | | | | |
| ^5^ Tail effective sample size | | | | | | | | | | |
