## Supplementary material for "Inferring a novel insecticide resistance metric and exposure variability in mosquito bioassays across Africa": S9 Table: S9_Table.html

|  |  |  |  |  |  |  |  |  |  |  |  |  |  |
| --- | --- | --- | --- | --- | --- | --- | --- | --- | --- | --- | --- | --- | --- |
| Model predictions per group | | | | | | | | | | | | | |
| from joint SB and EHT model | | | | | | | | | | | | | |
| Assay pair | ITN killing effect | | | LD_50_^1^ | | | μ_T_^2^ | | | σ_T_^3^ | | | Cor(μ_T_, σ_T_)^4^ |
|  | median | q025 | q975 | median | q025 | q975 | median | q025 | q975 | median | q025 | q975 |  |
| 1 | 3.51% | 1.81% | 5.67% | 2.60 × 10^2^ | 8.55 × 10^1^ | 2.01 × 10^3^ | 5.56 | 4.45 | 7.61 | 3.45 | 2.53 | 5.13 | 0.97 |
| 2 | 0.00% | 0.00% | 0.00% | 7.90 | 7.40 | 8.47 | 2.07 | 2.00 | 2.14 | 3.78 × 10^−1^ | 2.72 × 10^−1^ | 4.59 × 10^−1^ | 0.22 |
| 3 | 8.05% | 5.62% | 10.80% | 4.57 × 10^1^ | 2.79 × 10^1^ | 9.68 × 10^1^ | 3.82 | 3.33 | 4.57 | 3.22 | 2.63 | 4.11 | 0.93 |
| 4 | 9.27% | 6.12% | 12.80% | 1.73 × 10^1^ | 1.27 × 10^1^ | 2.69 × 10^1^ | 2.85 | 2.55 | 3.29 | 2.67 | 2.20 | 3.35 | 0.77 |
| 5 | 1.36% | 0.12% | 4.46% | 7.91 | 6.28 | 1.09 × 10^1^ | 2.07 | 1.84 | 2.39 | 1.24 | 9.01 × 10^−1^ | 1.69 | 0.60 |
| 6 | 0.00% | 0.00% | 0.00% | 7.57 | 7.21 | 7.94 | 2.02 | 1.98 | 2.07 | 4.91 × 10^−1^ | 4.00 × 10^−1^ | 5.83 × 10^−1^ | −0.02 |
| 7 | 6.99% | 4.38% | 9.86% | 4.23 | 3.84 | 4.67 | 1.44 | 1.35 | 1.54 | 1.44 | 1.31 | 1.59 | 0.20 |
| 8 | 42.09% | 25.15% | 56.92% | 6.39 × 10^−1^ | 1.97 × 10^−1^ | 9.39 × 10^−1^ | −4.48 × 10^−1^ | −1.62 | −6.32 × 10^−2^ | 1.24 | 4.84 × 10^−1^ | 8.18 | −0.50 |
| 9 | 27.53% | 25.89% | 29.21% | 1.79 × 10^2^ | 1.20 × 10^1^ | 8.98 × 10^3^ | 5.19 | 2.49 | 9.10 | 9.86 | 5.23 | 1.64 × 10^1^ | 0.99 |
| 10 | 15.59% | 12.53% | 18.70% | 1.86 × 10^3^ | 2.34 × 10^1^ | 1.51 × 10^6^ | 7.53 | 3.15 | 1.42 × 10^1^ | 8.15 | 3.75 | 1.46 × 10^1^ | 0.98 |
| 11 | 7.01% | 4.05% | 10.38% | 5.23 × 10^3^ | 2.11 × 10^1^ | 4.88 × 10^7^ | 8.56 | 3.05 | 1.77 × 10^1^ | 6.32 | 2.37 | 1.26 × 10^1^ | 0.98 |
| 12 | 86.37% | 82.36% | 90.02% | 4.77 × 10^−4^ | 5.78 × 10^−7^ | 3.05 × 10^−2^ | −7.65 | −1.44 × 10^1^ | −3.49 | 6.32 | 2.69 | 1.21 × 10^1^ | −0.97 |
| 13 | 40.85% | 37.32% | 44.47% | 4.65 | 1.41 | 3.58 × 10^1^ | 1.54 | 3.47 × 10^−1^ | 3.58 | 9.96 | 5.00 | 1.68 × 10^1^ | 0.79 |
| 14 | 28.78% | 21.99% | 35.56% | 5.57 × 10^1^ | 4.67 | 4.14 × 10^3^ | 4.02 | 1.54 | 8.33 | 8.61 | 4.00 | 1.55 × 10^1^ | 0.85 |
| 15 | 11.32% | 8.01% | 14.31% | 1.86 | 1.20 | 2.35 × 10^1^ | 6.21 × 10^−1^ | 1.86 × 10^−1^ | 3.16 | 1.09 | 5.67 × 10^−1^ | 3.43 | 0.98 |
| 16 | 11.73% | 8.46% | 14.96% | 1.10 × 10^3^ | 1.05 × 10^1^ | 2.19 × 10^6^ | 7.00 | 2.35 | 1.46 × 10^1^ | 6.50 | 2.46 | 1.30 × 10^1^ | 0.98 |
| 17 | 35.66% | 30.94% | 40.57% | 1.51 × 10^1^ | 2.53 | 3.18 × 10^2^ | 2.71 | 9.29 × 10^−1^ | 5.76 | 9.51 | 4.69 | 1.64 × 10^1^ | 0.83 |
| 18 | 13.69% | 5.14% | 21.85% | 7.41 × 10^3^ | 6.21 × 10^1^ | 1.76 × 10^7^ | 8.91 | 4.13 | 1.67 × 10^1^ | 8.82 | 4.16 | 1.53 × 10^1^ | 0.81 |
| 19 | 0.03% | 0.00% | 5.03% | 3.76 × 10^5^ | 3.34 × 10^1^ | 5.87 × 10^11^ | 1.28 × 10^1^ | 3.51 | 2.71 × 10^1^ | 3.66 | 1.81 × 10^−1^ | 1.07 × 10^1^ | 0.37 |
| 20 | 10.92% | 7.21% | 14.51% | 3.59 × 10^4^ | 1.77 × 10^2^ | 7.02 × 10^7^ | 1.05 × 10^1^ | 5.17 | 1.81 × 10^1^ | 9.10 | 4.67 | 1.52 × 10^1^ | 0.96 |
| 21 | 10.56% | 6.86% | 14.33% | 1.97 × 10^4^ | 1.01 × 10^2^ | 2.93 × 10^7^ | 9.89 | 4.62 | 1.72 × 10^1^ | 8.47 | 4.14 | 1.45 × 10^1^ | 0.96 |
| 22 | 72.53% | 65.50% | 78.76% | 6.34 × 10^−3^ | 4.75 × 10^−5^ | 1.07 × 10^−1^ | −5.06 | −9.95 | −2.23 | 7.32 | 3.13 | 1.40 × 10^1^ | −0.91 |
| 23 | 43.49% | 35.10% | 51.54% | 1.52 | 3.42 × 10^−1^ | 1.45 × 10^1^ | 4.21 × 10^−1^ | −1.07 | 2.67 | 7.60 | 3.01 | 1.47 × 10^1^ | 0.31 |
| 24 | 88.54% | 78.64% | 94.77% | 1.61 × 10^−1^ | 2.18 × 10^−5^ | 4.63 × 10^−1^ | −1.83 | −1.07 × 10^1^ | −7.69 × 10^−1^ | 9.02 × 10^−1^ | 8.17 × 10^−2^ | 7.36 | −0.98 |
| 25 | 25.74% | 20.66% | 31.01% | 1.21 × 10^2^ | 6.05 | 1.48 × 10^4^ | 4.79 | 1.80 | 9.60 | 8.52 | 3.84 | 1.52 × 10^1^ | 0.93 |
| 26 | 10.89% | 2.46% | 17.98% | 1.19 × 10^2^ | 1.94 | 5.58 × 10^5^ | 4.78 | 6.65 × 10^−1^ | 1.32 × 10^1^ | 4.62 | 7.09 × 10^−1^ | 1.19 × 10^1^ | 0.97 |
| 27 | 27.13% | 18.33% | 36.45% | 3.05 × 10^1^ | 2.76 | 3.28 × 10^3^ | 3.42 | 1.01 | 8.10 | 7.09 | 2.50 | 1.43 × 10^1^ | 0.83 |
| 28 | 7.79% | 7.00% | 8.56% | 7.01 × 10^−1^ | 5.61 × 10^−1^ | 8.14 × 10^−1^ | −3.55 × 10^−1^ | −5.77 × 10^−1^ | −2.06 × 10^−1^ | 1.68 × 10^−1^ | 6.64 × 10^−2^ | 2.82 × 10^−1^ | −0.43 |
| 29 | 4.31% | 3.43% | 5.20% | 7.32 × 10^−1^ | 5.92 × 10^−1^ | 8.54 × 10^−1^ | −3.12 × 10^−1^ | −5.25 × 10^−1^ | −1.58 × 10^−1^ | 1.46 × 10^−1^ | 2.44 × 10^−2^ | 2.54 × 10^−1^ | −0.46 |
| 30 | 4.43% | 1.60% | 7.04% | 1.01 | 9.41 × 10^−1^ | 1.13 | 1.43 × 10^−2^ | −6.06 × 10^−2^ | 1.19 × 10^−1^ | 3.76 × 10^−1^ | 2.06 × 10^−1^ | 6.03 × 10^−1^ | 0.38 |
| 31 | 0.48% | 0.00% | 4.56% | 7.96 × 10^−1^ | 6.25 × 10^−1^ | 9.32 × 10^−1^ | −2.28 × 10^−1^ | −4.70 × 10^−1^ | −7.01 × 10^−2^ | 5.67 × 10^−2^ | 2.46 × 10^−3^ | 1.71 × 10^−1^ | −0.30 |
| 32 | 7.03% | 0.00% | 17.93% | 9.56 × 10^−1^ | 8.24 × 10^−1^ | 1.16 | −4.45 × 10^−2^ | −1.93 × 10^−1^ | 1.52 × 10^−1^ | 3.89 × 10^−1^ | 5.45 × 10^−2^ | 8.31 × 10^−1^ | 0.28 |
| 33 | 66.79% | 61.72% | 71.45% | 4.54 × 10^−1^ | 3.14 × 10^−1^ | 5.98 × 10^−1^ | −7.90 × 10^−1^ | −1.16 | −5.14 × 10^−1^ | 1.48 × 10^−1^ | 8.95 × 10^−3^ | 3.45 × 10^−1^ | −0.46 |
| 34 | 88.41% | 83.24% | 92.46% | 3.61 × 10^−1^ | 1.98 × 10^−1^ | 5.33 × 10^−1^ | −1.02 | −1.62 | −6.29 × 10^−1^ | 2.13 × 10^−1^ | 1.41 × 10^−2^ | 5.11 × 10^−1^ | −0.75 |
| 35 | 49.02% | 44.41% | 53.70% | 6.40 × 10^−1^ | 1.66 × 10^−1^ | 2.63 | −4.46 × 10^−1^ | −1.80 | 9.66 × 10^−1^ | 1.06 × 10^1^ | 5.67 | 1.76 × 10^1^ | 0.05 |
|  |  |  |  |  |  |  |  |  |  |  |  |  |  |
| --- | --- | --- | --- | --- | --- | --- | --- | --- | --- | --- | --- | --- | --- |
| ^1^ Population median of lethal dose, equal to exp(μ_T_) (scale: multiples of discriminating dose) | | | | | | | | | | | | | |
| ^2^ Mean of log lethal dose (scale: log of multiples of discriminating dose) | | | | | | | | | | | | | |
| ^3^ Standard deviation of log lethal dose (scale: log of multiples of discriminating dose) | | | | | | | | | | | | | |
| ^4^ Pearson correlation coefficient | | | | | | | | | | | | | |
